# Loss of Zfp335 triggers cGAS/STING-dependent apoptosis of post-β selection pre-T cells

**DOI:** 10.1101/2021.12.03.471158

**Authors:** Jeremy J Ratiu, William Barclay, Qun Wang, Naren Mehta, Melissa J Harnois, Devon DiPalma, Sebastian Wellford, Sumedha Roy, Alejandra V Contreras, David Wiest, Yuan Zhuang

## Abstract

Production of a diverse peripheral T cell compartment requires massive expansion of the bone marrow progenitors that seed the thymus. There are two main phases of expansion during T cell development, following T lineage commitment at the DN2 stage and following successful rearrangement and selection for functional TCRβ chains in DN3 thymocytes, which promotes development of DN4 cells to the DP stage. Signals driving expansion of DN2 thymocytes are well studied, however, factors regulating the proliferation and survival of DN4 cells remain poorly understood. Here, we uncover an unexpected link between the transcription factor Zfp335 and control of cGAS/STING-dependent cell death in post-β-selection DN4 thymocytes. Zfp335 controls survival by sustaining expression of Ankle2, which suppresses cGAS/STING-dependent cell death. Together, this study identifies Zfp335 as a key transcription factor controlling the survival of proliferating post-β-selection thymocytes and demonstrates a key role for the cGAS/STING pathway driving apoptosis of developing T cells.

## Introduction

Development of large number of T cells with clonally acquired T cell receptor (TCR) in the thymus demands a small number of bone marrow derived progenitors to undergo vigorous expansion prior to each of the sequentially ordered TCR gene rearrangement events. The first major expansion occurs immediately upon T lineage commitment at the DN2 stage prior to rearrangement of any TCR gene^1–4^. The expanded T cell progenitors enter the DN3 stage where rearrangement at the TCRβ, γ, δ gene loci become permissive. In postnatal thymus, the majority of DN3 cells will choose the αβT cell fate due to the generation of a productively rearranged TCRβ chain. Post β-selection DN3 cells then move to the DN4 stage where the second phase of expansion occurs, typically involving several rounds of rapid proliferation over the course of 2-3 days in mice. The expansion of TCRβ positive cells result in generation of the post mitotic DP cells, which constitutes 90% of all thymocytes in post-natal mice and humans. DP cells undergo TCRα gene rearrangement and selection, a process resulting in approximately 1% of cells surviving and contributing to the peripheral T cell pool. Therefore, the expansion of post β-selection DN4 cells prior to TCRα gene rearrangement and TCR selection represents a critical amplifier to control the output of αβT cells from the thymus.

While most stages of T cell development have been subject to extensive genetic and functional characterization, the post-β-selection proliferative phase remains less well understood. Previous studies have shown that proliferation but not survival of DN4 cells is dependent upon IL-7R signaling which functions to repress Bcl6 expression^5^. Similarly, proliferation during this stage of development also requires the combined activities of NOTCH and pre-TCR signaling^6–9^. This effect is in part the result of induction of Fbxl1 and Fbxl12 which induce polyubiquitination and proteasomal degradation of Cdkn1b ensuring proper cell cycle progression and proliferation ^10^. Survival of proliferating post-β-selection thymocytes was found to require expression of the chromatin associated protein yin yang 1 (Yy1), the absence of which drives p53-dependent apoptosis^11^. Animal models exploring cell death during T cell development have repeatedly shown thymocyte apoptosis, including among DN4 cells, is largely driven by activities of pro-apoptotic Bcl2 family proteins^12–16^. Pathways controlling the survival and death of early proliferating thymocytes upstream of the Bcl2 family remain largely unexplored.

Underpinning the fate decisions of thymocytes are vast transcriptional networks which coordinate the intricate changes and checkpoint traversals required for proper development ^17^. Numerous transcription factors function at different stages to achieve this result. One transcription factors family of particular importance are the basic helix-loop-helix E proteins, which include E2A, HEB and E2-2. In developing T cells, activities of the E2A and HEB have been shown to regulate nearly all stages of thymopoiesis ^18, 19^. These E proteins play critical roles in enforcing the β-selection checkpoint by promoting expression of *Rag1/2* ^20^ and *pre-Tα* ^21^, activation of the TCRβ ^22^, TCRγ, and TCRδ loci ^23^ and preventing passage of DN cells lacking a functional TCRβ chain from progressing to the DP stage ^24, 25^. Additionally, E protein activity has been shown to enforce early T cell lineage commitment ^26^ and promote survival of post-β-selection DP thymocytes undergoing TCRα recombination ^27^. Together, the combined activities of E proteins play critical and indispensable roles in the establishment of a functional T cell repertoire. However, due to the widespread binding of these factors throughout the genome of developing thymocytes our understanding of their roles in development are far from complete.

The cGAS/STING pathway functions to sense cytosolic DNA and initiate innate immune responses ^28^. Cyclic GMP-AMP (cGAMP) synthase (cGAS) recognizes dsDNA, typically of foreign origin, catalyzing the generation of the cyclic dinucleotide (CDN) second messenger cGAMP which in turn drives STING activation and down-stream signaling ^29^. The cGAS/STING pathway is best known for its functions in non-immune and innate immune cells such as macrophage and dendritic cells in the context of viral or bacterial infections. In these contexts, activation of the pathway typically results in the production of type I interferons and other pro-inflammatory mediators. Recent work has shown that the cGAS/STING pathway is also highly active but functionally divergent within T cells, primarily driving type I interferon-independent responses and apoptosis ^30–33^. Under steady-state conditions the cGAS/STING pathway plays a minimal role in T cell development as evidenced by normal thymic T cell subset proportions and overall thymus size in cGAS or STING-deficient C57/BL6 mice ^32^. However, it remains to be determined whether the cGAS/STING pathway plays a role in sensing and responding to cell intrinsic stresses during thymic T cell development.

In this study we show that loss of Zinc finger transcription factor 335 (Zfp335), triggered cGAS/STING-mediated apoptosis among proliferating DN4 cells. Zfp335 was initially identified from genetic mapping of familial traits that cause a severe form of microcephaly ^34^. Using a conditional knockout mouse model ^34, 35^ we show that loss of Zfp335 promotes cGAS/STING dependent apoptosis among proliferating post-β-selection DN4 thymocytes, severe reduction in overall thymic cellularity and a near absence of peripheral T cells. Mechanistically, Zfp335 functions to suppress cGAS/STING activation through promoting Ankle2 expression which in turn regulates the cGAS inhibitor Baf^36^. The importance of cGAS/STING pathway among DN4 thymocytes was further demonstrated by their sensitivity to STING agonist and STING-mediated cell death in wild type mice. Thus, we have uncovered for the first time a role for the cGAS/STING pathway in regulating thymic T cell development and identify the Zfp335/Ankle2/Baf axis as the first transcriptional network functioning to regulate cGAS/STING activity.

## Results

### Zfp335, an E-protein target, is critical for T cell development

The E protein family of transcription factors are indispensable regulators of nearly every stage of T cell development^4, 17, 22, 24, 27, 37–40^. E proteins control complex transcriptional networks which remain incompletely understood. To gain deeper insight into mechanisms by which E proteins regulate T cell development, we previously performed E2A ChIP-seq to identify the genome-wide binding sites during T cell development^40^. We identified Zfp335 as an E protein target during T cell development (Fig S1A). Analysis of published data showed E protein-deficient thymocytes exhibit significantly reduced *Zfp335* expression (Fig S1B)^39^. Since germline deletion of Zfp335 is non-viable^34^ we utilized a conditional deletion model driven in which Cre expression is controlled by the E8III enhancer of *Cd8a* (E8III-cre) to allow functional assessment of Zfp335 in post-β-selection thymocytes^35^. There are conflicting reports regarding the deletion kinetics for this Cre^35, 41^, therefore, we began by assessing its activity across T cell development in our system (Fig S1C-D). Consistent with Dashtsoodol *et* al., we found E8III-cre is highly active immediately upon entry into DN3a with no recombination activity evident in the preceding DN2 stage. However, recombination does not appear to be complete until the DP stage.

We subsequently assessed *Zfp335^fl/fl^* E8III-cre (Zfp335cKO) mice for thymic T cell development. Deletion of *Zfp335* led to a significant reduction in total thymic cellularity (Fig 1A-B). This reduction in thymic cellularity is likely due to defects in the αβ lineage as numbers of γδ T cells were not altered (Fig 1C-D). Assessment of developmental stages revealed the reduction in thymocyte numbers of Zfp335cKO mice begins at the DN4 stage (Fig 1E-I).

**Figure 1.**
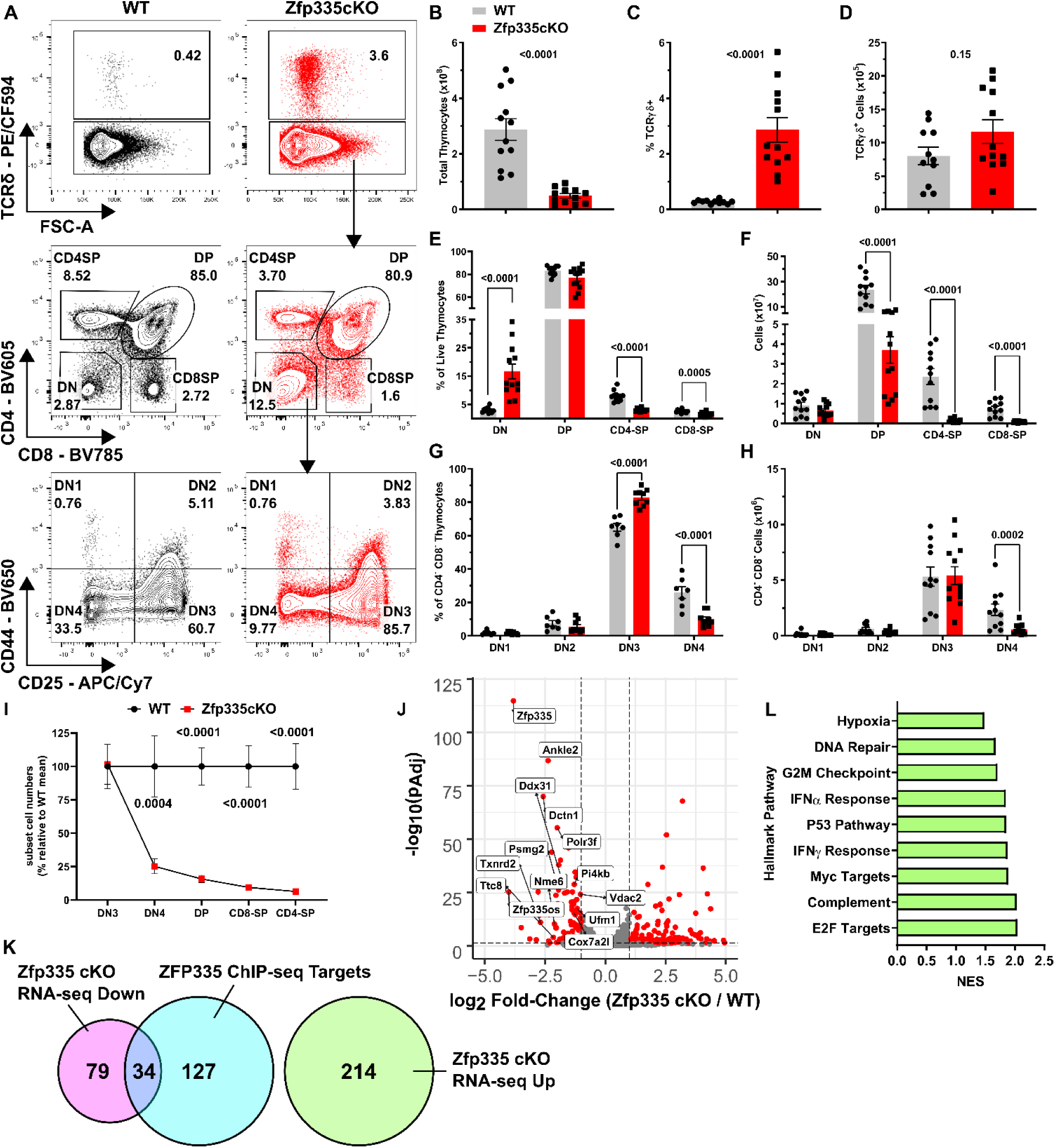
Zfp335 is critical to αβ T cell development. (A) Gating schema for *ex vivo* analysis thymocyte development beginning with live thymocytes (DAPI^-^ CD90.2^+^, gating not shown). (B) Total thymic cellularity in WT (Cre-negative) or *Zfp335^fl/fl^ E8_III_-cre* (Zfp335cKO) mice. Total numbers (C) and frequency (D) of TCRγδ^+^ cells in WT or Zfp335cKO thymuses. Numbers (E) and frequencies (F) of DN, DP, and SP thymocyte subsets in WT or Zfp335cKO thymuses. Numbers (G) and frequencies (H) of early DN1-DN4 thymocyte subsets in WT or Zfp335cKO thymuses. (I) Relative cells numbers in DN3-SP thymocyte subsets represented as percent of WT mean. (J) Differential expression of select Zfp335-target genes by RNA-seq. (K) Overlap between Zfp335 ChIP-seq (GSE58293) and differentially expressed genes in Zfp335cKO and WT DP. (L) Gene Set Enrichment Analysis of differentially expressed genes (K). Positive enrichment scores indicate pathways positively enriched in Zfp335cKO cells. (A-K) Cre-negative WT (n=11) and Zfp335cKO (n=12) 4-5-week-old male and female mice from four independent experiments. *P*-values determined by Two-way ANOVA with *post hoc* Sidak test. (I-K) RNA-seq analysis of *Zfp335^+/+^ E8_III_-cre* or Zfp335cKO DP thymocytes (n=3 each) of 6-week-old female mice from one experiment. Plots show mean ± sem.

Examination of the peripheral T cell compartment revealed significantly reduced numbers of splenic T cells in Zfp335cKO mice (Fig S2A-K). A previous study identified the hypomorphic *Zfp335^bloto^* allele as the causative mutation in a unique form of T lymphopenia^42^. Like *Zfp335^bloto^* mice, we found that peripheral T cells in Zfp335cKO mice were almost exclusively of an effector or memory phenotype suggesting these mice also exhibit a similar defect in the establishment of the naïve T cell compartment.

To determine the transcriptional changes resulting from loss of Zfp335 we performed RNA-seq on Zfp335cKO DP thymocytes. DP cells were used as they were the first population exhibiting complete deletion (Fig S1D). We found that loss of Zfp335 results in differential expression of 327 genes (113 down, 214 up; Fig 1K,J). Among the 161 Zfp335 ChIP-seq targets identified in thymocytes^42^, 34 were down-regulated in Zfp335cKO mice (Fig 1K). No Zfp335 target genes were up-regulated in Zfp335cKO samples (Fig 1K) corroborating previous findings that Zfp335 primarily functions as a transcriptional activator^34, 42^. Consistent with transcriptomic analyses of *Zfp335^bloto^* mice^42^, gene set enrichment analysis (GSEA) revealed significant enrichment among type I and type III interferon signaling and P53 signaling pathways in Zfp335cKO DP cells (Fig 1L). Together, these findings identify Zfp335 as a key transcription factor regulating T cell development.

### Loss of Zfp335 in DN3 thymocytes does not impair β-selection

Zfp335 deletion results in reduced cell numbers beginning at the DN4 stage, raising the possibility that the inability to rearrange the TCRβ locus could be responsible. Consequently, we assessed TCRβ rearrangement in DN3 and DN4 thymocytes by intracellular staining. The frequency of icTCRβ^+^ cells among Zfp335cKO DN3 and DN4 subsets was comparable to that of WT (Fig S3A-C). Therefore, TCRβ rearrangement and subsequent pre-TCR expression are unimpaired in Zfp335cKO mice.

In addition to pre-TCR expression, to successfully traverse the β-selection checkpoint, pre-TCR signals are required for release from cell cycle arrest, survival and progression to DP^43^. CD27 surface expression is increased by pre-TCR signals in DN3 thymocytes^44^. Zfp335cKO DN3 thymocytes exhibited CD27 upregulation comparable to that of WT (Fig S3D-E) indicating Zfp335-deficiency does not lead to impaired pre-TCR signaling. Together, these results indicate that the observed reduction of DN4 cells in Zfp335cKO mice did not result from failure to produce TCRβ subunits or failure to transduce pre-TCR signals.

### Zfp335 inhibits apoptosis during the DN-DP transition

Zfp335 deletion during the DN3 stage leads to severe defects in T cell development, likely during the post-β-selection proliferative phase. To determine if Zfp335-deficiency altered either the proliferation or survival of post-β-selection thymocytes, we directly measured these events in OP9-DL1 cultures *in vitro*^45^. Consistent with our *ex vivo* data, Zfp335cKO cells exhibit severely impaired progression to the DP stage (Fig 2A-B). Zfp335cKO cells exhibited modestly reduced proliferation compared to controls (Fig 2C-D). In contrast, Zfp335cKO cells underwent substantially increased rates of apoptosis (Fig 2E-F). Importantly, proliferation tracking (Fig 2G) and assessment of developmental progression (Fig 2H) of apoptotic mutant cells demonstrate they have undergone cell division and largely remain DN. These data suggest that Zfp335cKO cells are dying during the post-β-selection proliferative phase and that Zfp335 activity promotes the survival of DN4 thymocytes.

**Figure 2.**
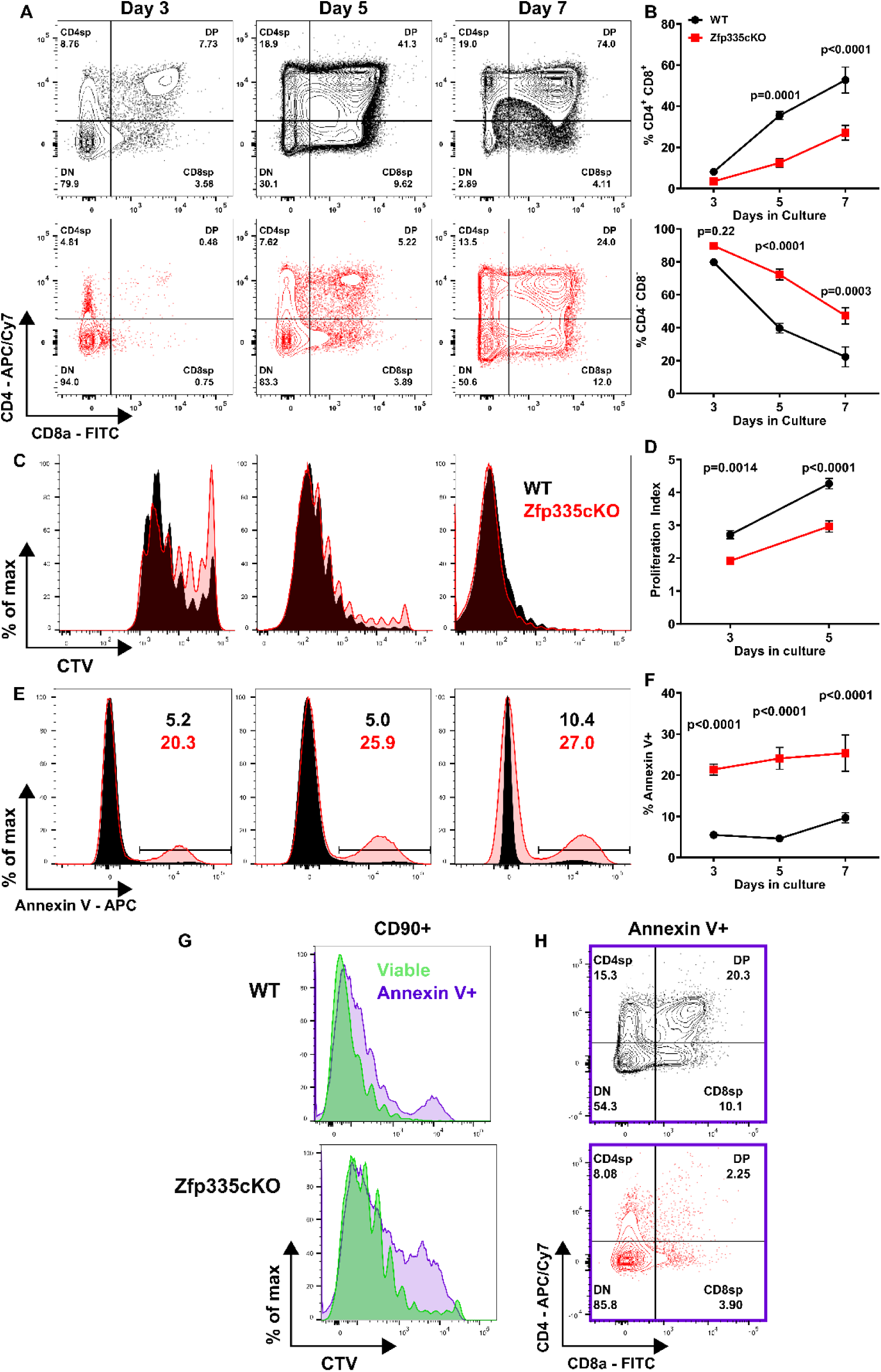
Zfp335cKO DN4 thymocytes undergo increased rates of apoptosis. (A-B) Assessment of developmental progression throughout OP9-DL1 culture. Proliferation assessment (C-D) by Cell Trace Violet (CTV) dilution and apoptosis analysis (E-F) based on Annexin V binding at day3, 5 or 7 of culture. (G) Representative comparison of CTV dilution between Annexin V^+^ and viable (DAPI^-^ Annexin V^-^) cells on day 5 of culture. Representative CD4 vs CD8 expression among Annexin V^+^ cells on day 5 of culture. n=6 WT or n=5 Zfp335cKO from three independent experiments. *P*-values determined using Two-way Repeated Meaures ANOVA with *post hoc* Sidak Test. Plots show mean ± sem.

### Ectopic Bcl2 expression rescues the developmental defect resulting from loss of Zfp335

Our RNA-seq studies revealed Zfp335cKO thymocytes exhibit increased expression of the pro-apoptotic Bcl2-family members, PUMA (*Bbc3*), NOXA (*Pmaip1*) and *Bax* (Fig 3A), suggesting that these factors may be responsible for the observed increase in apoptosis among Zfp335cKO thymocytes. The function of these proteins can be antagonized by ectopic expression Bcl2. Thus, we asked whether Bcl2 overexpression could rescue Zfp335cKO thymocyte apoptosis. WT or Zfp335cKO DN3/4 thymocytes were transduced with control or Bcl2-expressing retroviruses then grown in the OP9-DL1 culture system. Bcl2 overexpression significantly reduced apoptosis in Zfp335cKO cells, indicating the induction of pro-apoptotic Bcl2 family members was at least partially responsible for the observed increase in apoptosis in Zfp335-deficient thymocytes (Fig 3B-C).

**Figure 3.**
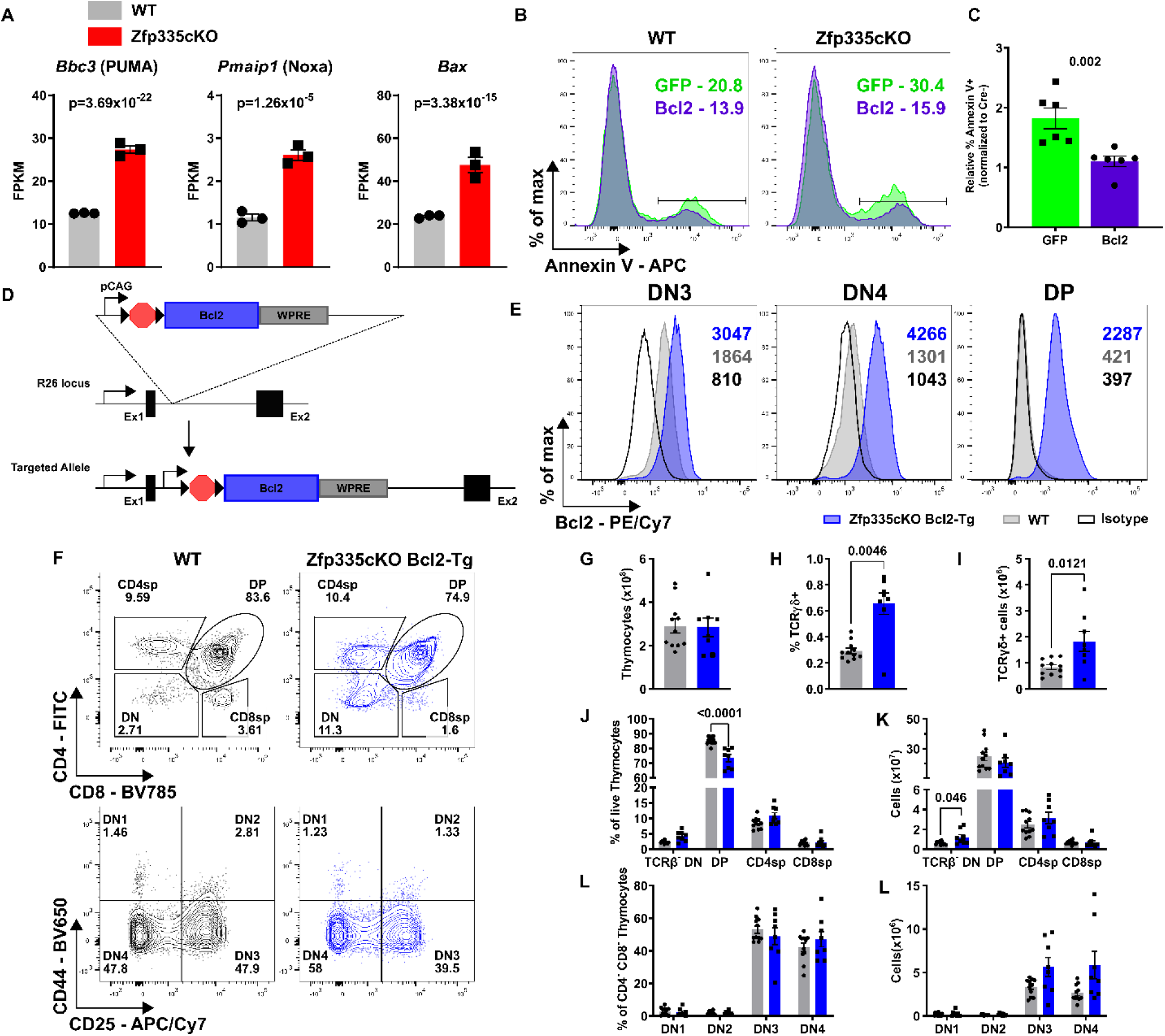
Bcl2 overexpression rescues Zfp335-deficient thymocytes from apoptosis. (A) Expression of pro-apoptotic Bcl2 family genes *Bbc3*, *Pmaip1*, or *Bax* from RNA-seq of control or *Zfp335^fl/fl^ E8_III_-cre* DP thymocytes. Representative gating (B) and quantification of apoptosis among *Zfp335^fl/fl^ E8_III_-cre* thymocytes transduced with Bcl2 or GFP RV after 5 days of OP9-DL1 culture (n=5). (E) Representative expression of isotype control (open black) or Bcl2 in WT (grey) or *Zfp335^fl/fl^ R26^LSL-Bcl2^ E8_III_-cre* (blue) DN3, DN4 or DP thymocytes. (F) Gating for identification of thymocyte subsets in WT WT (grey) or *Zfp335^fl/fl^ R26^LSL-Bcl2^ E8_III_-cre* (blue) mice. DN1-4 gating pre-gated on TCRβ^-^. (G) Total thymocyte numbers. Total numbers (H) and proportions (I) of TCRδ^+^ cells. Frequencies (J) and total numbers (K) of DN, DP, CD4-SP and CD8-SP thymocytes. Frequencies (L) and total numbers (M) of DN1-DN4 thymocytes. (F-M) n=11 WT or n=8 *Zfp335^fl/fl^ R26^LSL-Bcl2^ E8_III_-cre.* Data compiled from one (A), two (B-C) or five (D-L) independent experiments. P-values determined by Wald test (A), Mann-Whitney U-test (C) or Two-way ANOVA with *post hoc* Sidak’s test (H-M). Plots show mean ± sem.

We next sought to test the ability of Bcl2 overexpression to rescue Zfp335-deficient cells from apoptosis *in vivo* through generating Bcl2 conditional transgenic mice (Fig 3D). Intracellular staining revealed that *Zfp335^fl/fl^ R26^LSL-Bcl2-Tg^* E8_III_-cre (Zfp335cKO Bcl2-Tg) thymocytes exhibited increased Bcl2 protein expression relative to WT (Fig 4E). Phenotypic analysis demonstrated that ectopic Bcl2 expression was able to fully rescue the early developmental defects observed in Zfp335-deficient mice, restoring traversal of the β-selection checkpoint, transition to the DP stage, and total thymic cellularity (Fig 3F-L).

**Figure 4.**
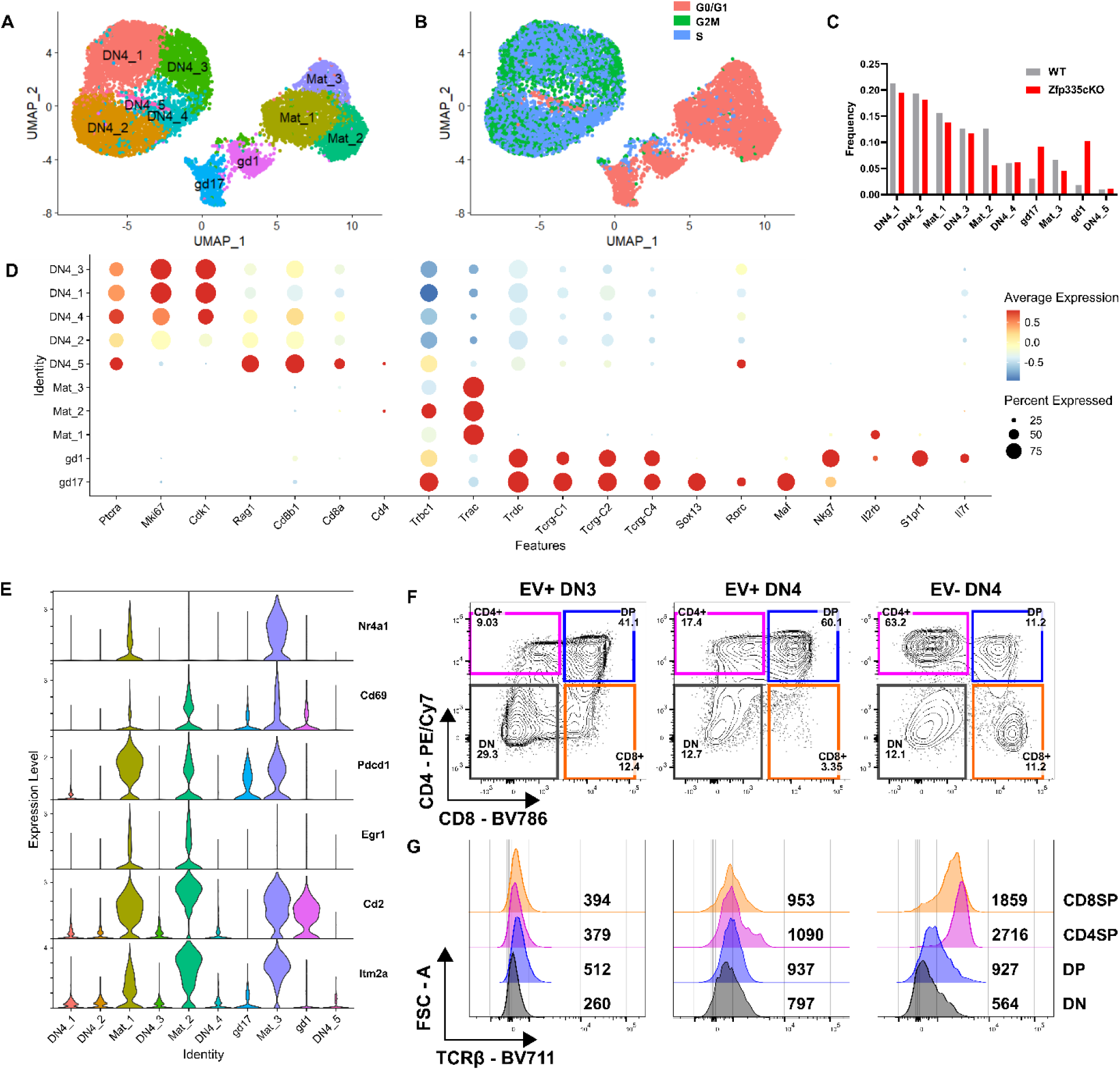
Defining the ‘true’ DN4 thymocyte population at the single cell level. (A) UMAP projection and identification of 10 clusters identified in full scRNA-seq dataset. (B) UMAP colored by cell cycle phase. Blue or green identify actively cycling cells. (C) Frequency distributions for WT (n=6357) and Zfp335cKO (n=5392) cells across the ten clusters. (D) Dot plot of key cell type-defining genes. (E) Violin plots of positive selection signature genes in thymocytes (Mingueneau et al. 2013). (F) Representative gating for CD4 vs. CD8 expression on day 3 of OP9-DL1 cultures seeded with WT Thy1.1 retrovirus transduced (EV+) DN3 or DN4 cells or non-transduced (EV-) DN4 cells. (G) Representative TCRβ expression among DN, DP, CD4SP or CD8SP cells from (F). Numbers indicate geometric MFI of TCRβ expression. (F-G) Data representative of two independent experiments.

Consistent with studies of *Zfp335^bloto^* mice^42^, Bcl2 overexpression failed to rescue the impairment in final single positive thymocyte maturation (Fig S4A-C) or peripheral T cell compartment numbers (Fig S4D-E) and effector status (Fig S4F-H). Taken together, these data suggest that the early impairment of thymocyte development following loss of Zfp335 expression is due to increased rates of DN4 apoptosis driven by pro-apoptotic Bcl2-family members. However, our *in vivo* studies also revealed an additional, Bcl2-independent late block in terminal T cell differentiation within the thymus.

### Defining the ‘true’ DN4 thymocyte population at the single cell level

The DN4 stage of T cell development remains poorly understood and, as a result, poorly defined. DN4 cells are identified by lack of expression of identifying markers associated with any other thymocyte subset. Based on these criteria, it is possible that DN4 cells defined by marker exclusion may not be homogenous. To assess whether there is any heterogeneity in the DN4 compartment exacerbated by Zfp335-deficiency, we performed scRNA-seq of phenotypically defined DN4 cells. After quality control, libraries yielded transcriptome data for 6,537 or 5,392 high-quality cells from WT or Zfp335cKO samples, respectively.

We identified 10 unique cell clusters (Fig 4A-C). Five clusters were largely cycling cells (DN4_1-5; Fig 4A-B) uniquely expressing *Ptcra* (pre-Tα) and proliferation associated genes (*Mki67, Cdk1*) (Fig 4D), representing bona fide DN4 cells. Three clusters (Mat_1-3) expressed high levels of *Trac and Trbc1* transcripts (Fig 4D). Two additional clusters (gd17 and gd1) of γδ T cells were identified. gd17 cells express high levels of *Sox13*, *Rorc* and *Maf*, features of γδ17 while gd1 express *Nkg7*, *Il2rb*, *S1pr1* and *Il7r* associated with cytotoxic γδ T cells (Fig 4D). Based on this clustering, Zfp335-deficiency led to substantial proportional increases and decreases in the γδ T cell clusters and Mat_2 cluster relative to WT control, respectively (Fig 4C).

We were surprised to find a large proportion of phenotypically defined DN4 thymocytes expressing *Trac* transcripts and sought to define these populations. Consistent with their lack of surface CD4 or CD8 these cells uniformly lacked *Cd4, Cd8a* and *Cd8b1* transcripts (Fig 4D). We hypothesized that these cells may represent post-positive selection thymocytes that transiently down-regulated surface TCR, CD4 and CD8 expression. Consistent with our hypothesis, we found these cells express high levels of *Nr4a1*, *Cd69*, *Pdcd1*, *Egr1*, *Cd2*, and *Itm2a,* signature genes of positive selection ^46^. Based on this profile we define cells from these clusters as maturing αβ T cells.

Importantly, most cells associated with the maturing αβ or γδ T cell clusters were non-cycling (Fig 4B), and therefore, not ‘true’ DN4 cells. Retroviral transduction depends on cell being cycling^47^. Therefore, we determined whether ‘true’ DN4 cells could be separated from contaminating populations *ex vivo* with retroviruses. Virally transduced or non-transduced DN4 cells were placed in OP9-DL1 culture. Non-transduced DN4 cells preferentially give rise to single-positive cells expressing high levels of surface TCR, whereas, transduced DN4 become DP (Fig 4F-G). Since OP9-DL1 cells are unable to support positive selection, we conclude that these non-transduced DN4 cells are post-positive selection cells transitioning to SP. Together, these results demonstrate that the phenotypically defined DN4 compartment is heterogenous and establishes retroviral transduction as a method to isolate DN4 cells for *in vitro* analysis.

### Ankle2 is a critical Zfp335-regulated gene required for survival of DN4 thymocytes

Next, we focused our scRNA-seq analyses on determining the transcriptional changes in DN4 cells resulting from loss of Zfp335. Maturing αβ and γδ cells were removed leaving only ‘true’ DN4 cells. Based on recombination kinetics (Fig S1D) not all Zfp335cKO DN4 cells have undergone deletion. *Zfp335* expression could not reliably delineate mutant from non-mutant cells due to low detection rate (8% of Zfp335cKO vs 17.7% of WT cells). To identify true mutant DN4 cells in our dataset, we assessed transcription factor activity using gene set scores calculated for each cell based on the expression of the Zfp335 ChIP-seq target genes down-regulated in mutant DP cells (Fig 1J,K). Zfp335cKO cells exhibited a bimodal distribution for the gene set. Using established methods^48^, cutoff values were determined for the distribution and cells falling below this threshold were considered true mutants (Fig S5A). Cutoffs were confirmed by differential expression analysis between WT and Zfp335cKO targets high or Zfp335cKO targets low cells. Compared to WT, Zfp335cKO targets low cells exhibited differential expression of 80 genes (60 down, 20 up; Fig S4B) whereas Zfp335cKO targets high cells only exhibited differential expression of 7 genes (5 down, 2 up; Fig S4C).

Zfp335cKO cells above the threshold were considered non-mutant, removed and the remaining cells were then reanalyzed (Fig 5A) identifying 8 unique clusters (Fig S4D). WT and mutant cells were distributed across each cluster. C1-3 were enriched for WT whereas C4 was almost entirely mutant cells (Fig S4E). Despite regression of standard cell cycle-associated genes, clustering was largely dictated by cell cycle (Fig S4F-I). We observed no differences in cell cycle phase distributions between WT and mutant (Fig S4H). Therefore, we chose to compare WT and mutant DN4 cells based on genotype. Among the 60 down-regulated genes in mutant DN4 cells, 44 are Zfp335 targets by ChIP-seq (Fig 5B)^42^. We hypothesized that reduced expression of one or more of these genes was responsible for the increased rates of apoptosis observed in mutant DN4 cells. Thus, we examined expression of the 12 Zfp335 target genes with experimental evidence demonstrating a negative regulatory role in cell death (Fig 5C-D). Four exhibited reduced expression in mutant DN4 thymocytes (Fig 5C). Examination of expression frequency identified *Ankle2* to have the greatest reduction in percent of mutant cells expression (Fig 5E).

**Figure 5.**
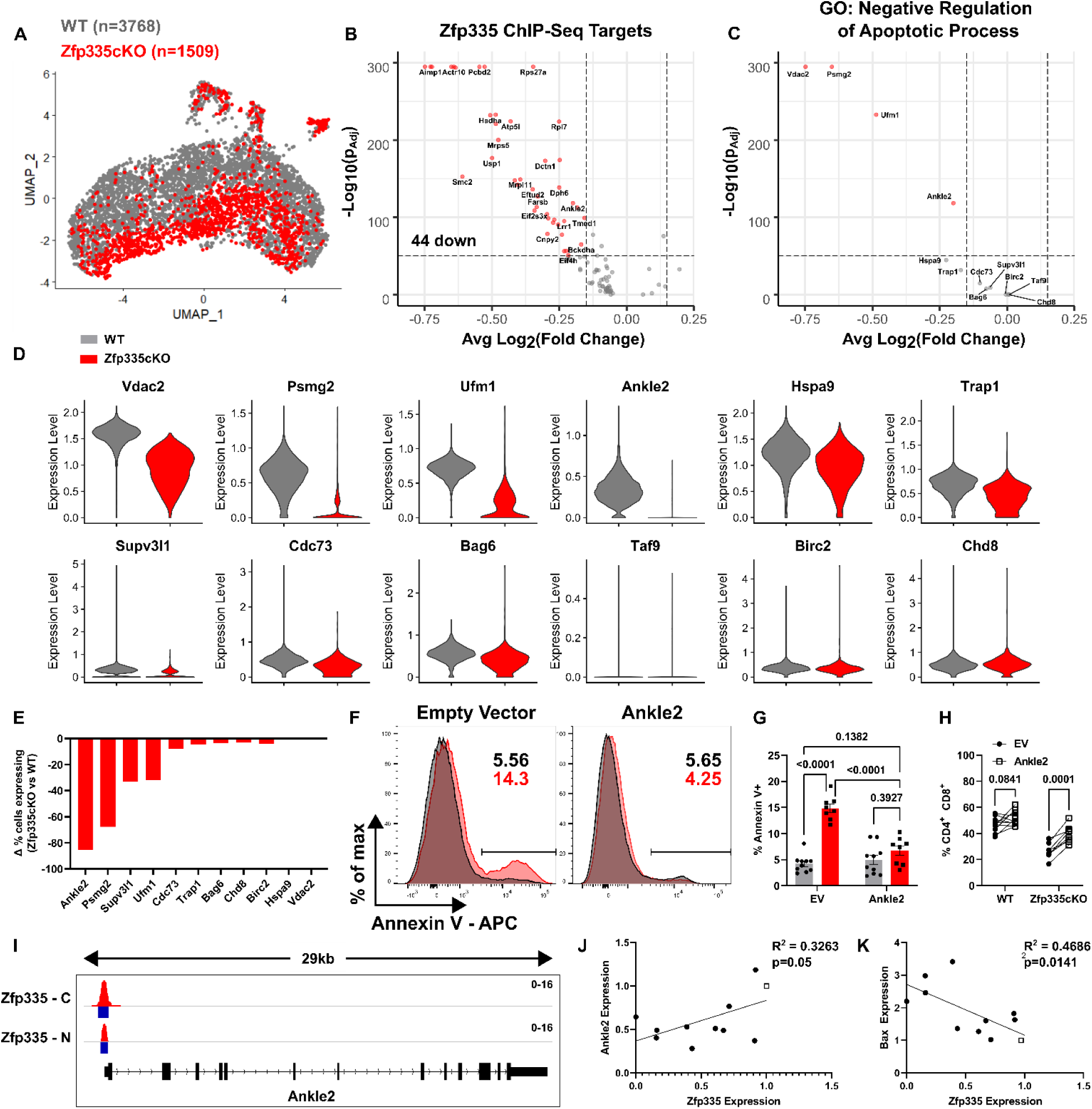
scRNA-seq identifies Ankle2 as a critical Zfp335-regulated gene controlling survival of DN4 thymocytes. (A) Violin plot of gene set score for Zfp335 target genes down-regulated in mutant DP thymocytes (Fig 1L-M) and cutoff value used to identify true Zfp335 mutant cells (black box). (B) UMAP projections colored by genotype. Volcano plot of all differentially expressed Zfp335 target genes (C) or those experimentally shown negatively regulate apoptotic processes (D) between Zfp335 mutant and WT cells. (E) Violin plots of anti-apoptotic Zfp335 target gene expression between Zfp335 mutant and WT DN4 cells. (F) Differential proportions of Zfp335 mutant cells expressing anti-apoptotic genes from E compared to WT cells. Representative gating (F) and quantification of apoptosis (G) or DP cell frequency (H) for EV or Ankle2 retrovirus transduced WT (n=10) or Zfp335cKO (n=8) DN3 thymocytes cultured on OP9-DL1 cells for 3 days. (I) Zfp335 ChIP-seq track of *Ankle2* locus in WT thymocytes (Zfp335-C or Zfp335-N antibodies, GSE58293). Blue boxes indicate significant binding peaks. Correlation between Ankle2 (J) or Bax (K) and Zfp335 expression in Scid.adh.2c2.SunTag CRISPRi cells expressing non-targeting (open squares) or Zfp335-targeting (closed circles) gRNAs. Data are compiled from one (A-E), two (J-K) or three (F-H) independent experiments. *P*-values determined by Wilcoxon Rank Sum test (B-C), two-way ANOVA with *post hoc* Tukey’s test for multiple comparisons (G), repeated measures ANOVA with Sidak’s test (H) or simple linear regression (J-K). Plots show mean ± sem.

*Ankle2* encodes an ER-restricted ankyrin repeat and LEM domain-containing protein^49^. *Ankle2* was recently identified as a critical Zfp335-regulated factor in the establishment of the naïve T cell^42^. Therefore, we tested whether *Ankle2* overexpression could rescue Zfp335cKO apoptosis. WT or Zfp335cKO DN3 thymocytes were transduced with EV or Ankle2 retrovirus and cultured on OP9-DL1 cells. Importantly, *Ankle2* overexpression was able to fully rescue Zfp335-deficient thymocytes from increased rates of apoptosis (Fig 5F-G). Moreover, *Ankle2* overexpression led to significantly increased proportions of DP cells among Zfp335cKO samples (Fig 5H).

Next, we sought to confirm that Ankle2 expression is directly regulated by Zfp335 in pre-T cells. Analysis of published ChIP-seq data showed Zfp335 binds the proximal promoter of *Ankle2* in thymocytes (Fig 5I). To examine the relationship between *Zfp335* and *Ankle2* expression we utilize the DN4-like mouse thymocyte cell line *Scid.adh.2c2*^50^ for CRISPR-based transcriptional inhibition (CRISPRi) studies^51^. These cells were transduced with retroviruses expressing *Zfp335* promoter-targeting gRNA and anti-GCN4scFv-sfGFP-KRAB fusion construct. *Zfp335*-targeted cells exhibited reduced *Ankle2* expression proportional to the efficiency of *Zfp335* knock-down (KD) (Fig 5J). Additionally, *Zfp335*KD resulted in increased expression of *Bax* like that observed in Zfp335cKO thymocytes (Fig. 5K). Together, these results demonstrate a direct relationship between *Zfp335* and *Ankle2* expression in developing T cells and suggest reduced *Ankle2* expression resulting from loss of Zfp335 drives DN4 apoptosis in Zfp335cKO mice.

### Disruption of the Zfp335/Ankle2/Baf axis drives cGAS/STING-dependent apoptosis of DN4 thymocytes

Next, we sought to determine the mechanism driving this increase in cell death resulting from reduced *Ankle2* expression. Ankle2 has previously been shown to control nuclear envelope (NE) reassembly and integrity following mitosis through regulation of Barrier to Autointegration Factor 1 (Banf1 or Baf) phosphorylation. Consistent with reduced *Ankle2* expression we observed significant increases in Baf phosphorylation among Zfp335cKO DN4 thymocytes (Fig 6A-C). Additionally, as previously reported, disruption of *ANKLE2* or *BANF1* expression in Hela cells led to severe disruptions in NE architecture (Fig S6A). To determine if the same is true for Zfp335-deficient DN4 thymocytes we examined the NE *ex vivo*. Indeed, Zfp335cKO DN4 thymocytes exhibit significantly altered NE architecture characterized by diffuse Lamin B1 throughout the cytosol, reduced DAPI signal possibly the result of loss of nuclear-cytosolic compartmentalization, and reduced nuclear sphericity (Fig 6D-G). Together, these data confirm that loss of Zfp335 leads to significantly altered NE architecture consistent with dysregulation of Ankle2/Baf-mediated NE reassembly and maintenance.

**Figure 6.**
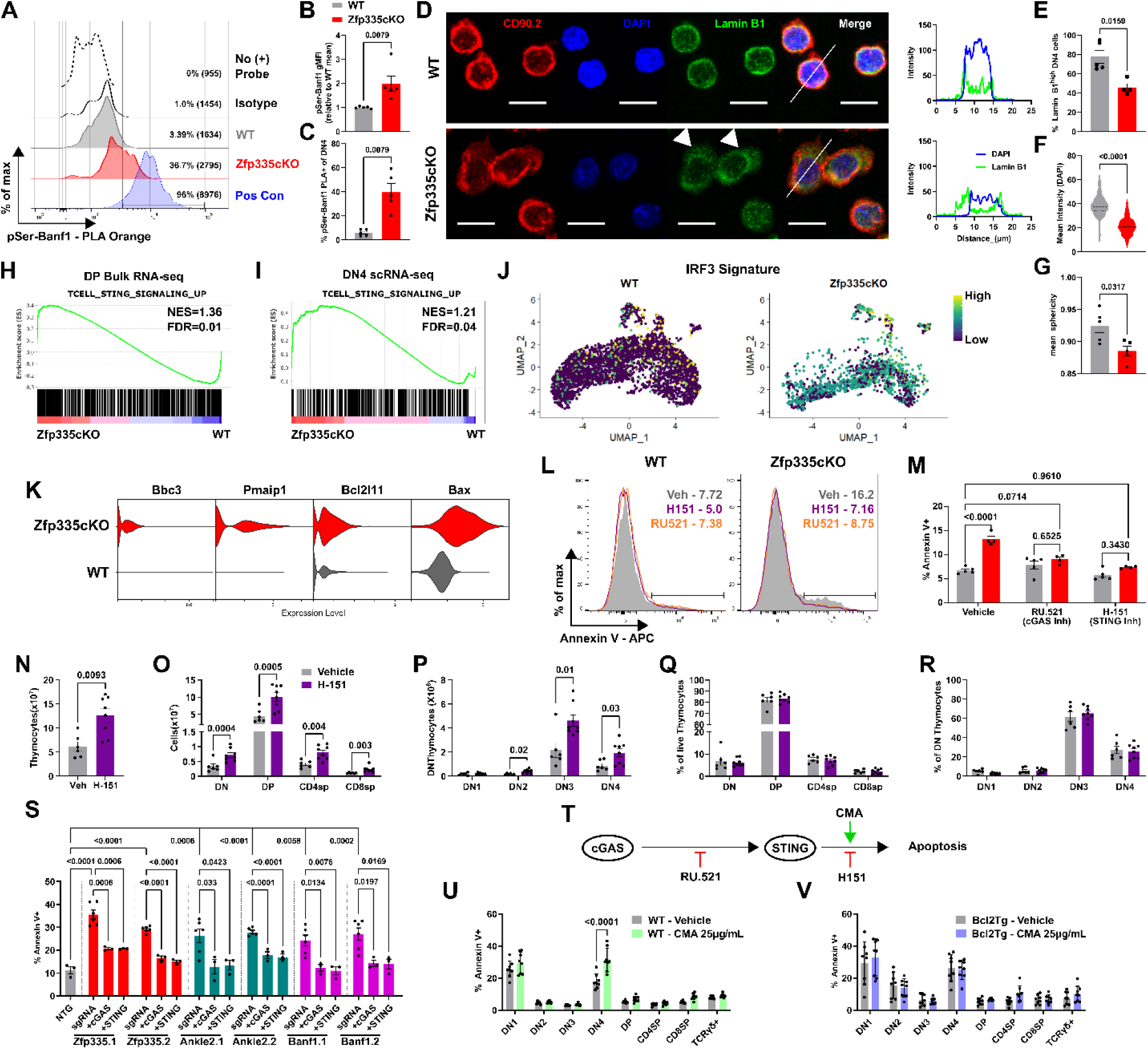
The Zfp335/Ankle2/Baf axis suppresses cGAS/STING-mediated apoptosis of DN4 thymocytes. (A) Representative histograms and gating of Baf phosphorylation as measured by proximity ligation assay (PLA). Percent phosphoserine-Baf and geometric MFI in parenthises are shown. Phosphoserine-Lamin B1 PLA was used as positive control. Quantification of Baf phosphorylation based on geometric MFI (B) or percent positive cells (C). (D) Representative immunofluorescence images of full cell thickness maximum intensity projections (left) and profile plots (right) of nuclear envelope staining in *ex vivo* DN4 thymocytes. Profile plots are based on white lines shown in merged images. Scale bars represent 10µm. Quantification of frequency of cells with high nuclear-associated Lamin B1 (E), mean DAPI pixel intensity (F) or mean nucleus sphericity (G) for *ex vivo* DN4 thymocytes. GSEA enrichment plots for T cell-specific STING signaling gene signature in DP bulk (H) or DN4 scRNA-seq data sets (I). (J) UMAP projection of IRF3 gene signature in WT or Zfp335 mutant DN4 thymocytes. (K) Violin plots of pro-apoptotic Bcl2 gene expression in WT or Zfp335 mutant DN4 thymocytes. Representative histograms (L) and quantification (M) of Annexin V-binding for WT or Zfp335cKO DN4 thymoctes treated with cGAS (RU.521) or STING (H-151) inhibitors or vehicle control and cultured on OP9-DL1 stromal cells for three days. Total thymocyte (N), DN, DP, CD4SP and CD8SP or DN1-DN4 cell numbers (O,P) or frequencies (Q,R) for Zfp335cKO mice treated with H-151 or vehicle *in vivo* for 7 days. (I-J) Thymocyte subset proportions for H-151 or vehicle treated Zfp335cKO mice. (S) Quantification of Annexin V binding among DN4 cells from *R26^LSL-Cas9^ Tcrd^CreERT2^* thymocytes transduced with gRNA-expressing retroviruses and cultured for three days on OP9-DL1 cells with 4-hydroxytamoxifen. (T) Schematic diagram of inhibitors (RU.521 or H-151) or agonists (CMA) used to study cGAS/STING-dependent apoptosis of DN4 thymocytes. Percent apoptosis induced by small molecule activation of STING among WT (U) or Zfp335cKO Bcl2Tg (V) thymocyte subsets. Values calculated by subtracting % Annexin V+ of vehicle-treated from % Annexin V+ of STING agonist-treated for each sample. *P*-values determined by Mann Whitney U-test (B-G,N) or two-way ANOVA with *post hoc* Tukey’s test (M) or Sidak’s test (O-P,U,V) or one-way ANOVA with post hoc Tukey’s test (S). Data shown are compiled from one (H-K), two (L-M), three (A-G,U,V) or five (N-R) independent experiments. Plots show mean ± sem or mean and interquartile range (F).

Accumulation of cytosolic DNA or exposure of nuclear contents to the cytosol via NE disruption have been shown to activate the cGAS/STING pathway^36, 52^. In T cells, cGAS/STING signaling generally results in anti-proliferative and pro-apoptotic effects^30, 31, 33, 53^. Therefore, we hypothesized that NE defects resulting from disruption of the Ankle2-Banf1 pathway downstream of Zfp335 loss drives cGAS/STING activation. Consistent with this hypothesis, GSEA revealed an enrichment for genes upregulated by T cells in response to STING signaling in both our bulk DP and single-cell DN4 datasets (Fig 6H,I). Additionally, we found increased IRF3 activity among mutant cells (Fig 6J). cGAS/STING-mediated death of mature T cells occur in part, due to increased expression of pro-apoptotic Bcl2 family genes^31^. Like our findings from bulk RNA-seq (Fig 4A), we also observed increased expression of Bbc3 (PUMA), Pmaip1 (NOXA), Bcl2l11 (Bid) and Bax among Zfp335cKO DN4 cells in our scRNA-seq dataset (Fig 6K).

In addition to nuclear DNA, mitochondrial DNA (mtDNA) serves as a substrate for cGAS^54^. mtDNA release requires mitochondrial outer membrane permeabilization resulting in mitochondrial membrane depolarization^55^. Examination of mitochondria showed Zfp335cKO thymocytes exhibit normal mitochondrial membrane potential and total mitochondrial mass (Fig S6B-C). Therefore, mtDNA release is unlikely to be driving cGAS/STING-mediated death following loss of Zfp335. Instead, exposure of gDNA to cytosolic cGAS resulting from disrupted nuclear envelope architecture is the most likely cause.

To test the contribution of cGAS/STING to increased rates of DN4 apoptosis in Zfp335cKO mice ‘true’ DN4 cells were isolated by EV viral transduction then placed in OP9-DL1 culture for 3 days with cGAS (RU.521)^56^ or STING (H-151)^57^ inhibitors. Chemical inhibition of either cGAS or STING fully rescued Zfp335cKO DN4 cells from death (Fig 6L,M). Additionally, Zfp335cKO mice receiving H-151 for 7 days exhibited significantly increased numbers of total thymocytes compared to vehicle controls (Fig 6N). Importantly, this increase in cellularity was primarily due to increased DP numbers (Fig 6O-R). Due to the short duration of treatment, we conclude that the increase in DP cells among H-151-treated Zfp335cKO mice is the result of reduced cell death during the preceding proliferative DN4 stage.

Next, we sought to determine the role of the Zfp335/Ankle2/Baf axis in suppressing the cGAS/STING-mediated apoptosis in DN4 cells. To test this, *R26^LSL-Cas9^ Tcrd^CreERT2^* DN3/DN4 thymocytes^58^ were transduced with retroviruses expressing *Zfp335, Ankle2, or Banf1* (encoding Baf) and *Mb21d1* (encoding cGAS) or *Tmem173* (encoding STING)-targeting gRNAs or non-targeting control gRNAs (NTG) then cultured for three days with OP9-DL1 cells in the presence of 4-hydroxytamoxifen. Consistent with conditional deletion, Cas9 targeting of *Zfp335* lead to a substantial increase in DN4 apoptosis (Fig 6S). Additionally, targeting of *Ankle2* or *Banf1* similarly lead to increased DN4 apoptosis. Importantly, these increases in apoptosis were cGAS/STING-dependent (Fig 6S). Similar results were observed when Cas9 expression was controlled by E8_III_-cre (Fig S6D-E). Together, these results demonstrate that disruption of the Zfp335/Ankle2/Baf axis drives cGAS/STING-mediated apoptosis of post-β-selection DN4 thymocytes.

### DN4 thymocytes are uniquely sensitive to cGAS/STING-mediated cell death

Finally, we sought to determine whether sensitivity to cGAS/STING-driven cell death is a unique feature of Zfp335cKO DN4 cells or a mechanism of the DN4 stage. DN-enriched WT thymocytes were treated with the STING agonist cridanimod (CMA) overnight then assayed for apoptosis. Interestingly, we found DN4 cells are uniquely sensitive to STING-mediated apoptosis (Fig 6T-U). Additionally, viability of Zfp335cKO Bcl2-Tg thymocytes was not impacted by CMA treatment (Fig 6V) suggesting that induction of pro-apoptotic Bcl2 family members downstream of STING activation are necessary for apoptosis of DN4 thymocytes.

Together, these data demonstrate that activation of the cGAS/STING pathway is a major contributor to Zfp335cKO DN4 apoptosis and that WT DN4 cells are uniquely sensitive to cGAS/STING-mediated death. Altogether, our studies demonstrate that loss of Zfp335 leads to defective T cell development resulting from dysregulation of the Zfp335/Ankle2/Baf axis ultimately driving cGAS/STING-mediated DN4 cell death.

## Discussion

In this study, we identify Zfp335 as a critical transcription factor regulating early T cell development within the thymus. Specifically, it functions to promote survival of proliferating cells following β-selection. Conditional deletion of Zfp335 led to severe reductions in all T cell populations beginning at the DN4 stage of development. Mechanistically, we show that reduced expression of the Zfp335-regulated gene Ankle2 is responsible for increased sensitivity to cell death and disruption of the Zfp335/Ankle2/Baf pathway controlling NE architecture drives cGAS/STING-dependent DN4 apoptosis.

Our studies provide the first comprehensive assessment of the heterogeneity within the DN4 thymocyte compartment at the single cell level. Surprisingly, phenotypically defined DN4 cells consist of cycling cells expressing pre-Tα which represent ‘true’ DN4 cells and mature or maturing αβ and γδ T cells. Positive selection of DP thymocytes induces a slight, transient down-regulation of CD4 and CD8^59^, however, the maturing αβ cells identified in our dataset completely lack both protein and mRNA expression. The cells we identified expressing TCRα transcripts exhibited expression patterns consistent with positive selection^46^ and therefore, are likely post positive-selection cells which have transiently lost surface expression of TCR, CD4 and CD8. Alternatively, these cells may have undergone positive selection without ever expressing CD4 or CD8. Regardless, these maturing cells may represent a novel developmental path within the thymus. However, more detailed studies will be needed to fully characterize these cells and determine if they represent a unique lineage or simply a rare differentiation path that can be taken by any positively selected cell.

Han *et al.* recently identified a hypomorph allele of *Zfp335* (*Zfp335^bloto^*) as the causative mutation leading to reduced total peripheral T cells and an almost complete absence of naïve T cells^42^. They found Ankle2 to be a critical Zfp335-regulated gene controlling late stages of thymic T cell maturation. However, the mechanism by which Ankle2 regulates maturation, and the establishment of the naïve T cell compartment remains unclear. The lack of apparent developmental defects in *Zfp335^blt/blt^* mice during early T cell development is likely due to their use of a hypomorph allele instead of a conditional knock out as *Zfp335^blt/blt^* mice exhibited normal expression of Ankle2 during the DN4 stage.

We have shown that Zfp335 is at least partially regulated by E protein activity in developing T cells. E proteins play numerous indispensable roles throughout organismal development, including T cell development ^4, 22, 37–40, 60–62^. However, due to widespread binding throughout the genome, the roles for transcriptional networks established by E proteins remain incompletely understood^40^. Our studies identify Zfp335 as a novel transcription factor downstream of E proteins critical to T cell development.

To date, studies of T cell-intrinsic roles for cGAS/STING pathway have largely focused on activation via synthetic STING agonists^31, 33, 53^ or expression of constitutive gain-of-function STING mutations^30^. These studies have primarily focused on roles of this pathway in mature peripheral T cells. To our knowledge, this is the first report of a physiological role for cGAS/STING in T cell development. Additionally, our identification of the Zfp335/Ankle2/Baf axis as key in repression of cGAS is the first transcriptional pathway identified which functions to prevent cGAS activation by self-DNA.

Baf was recently identified as a key inhibitor of cGAS sensing of self-DNA through competitive binding^36^. The ability of Baf to bind DNA is dependent upon its dephosphorylation which has been shown to be controlled by Ankle2 during mitotic exit^49^. Therefore, we propose the following mechanism by which loss of Zfp335 drives cGAS/STING-mediated apoptosis of DN4 thymocytes. Loss of Zfp335 results in impaired Ankle2 expression which in turn leads to the failure of Baf dephosphorylation during division. Baf hyperphosphorylation leads to improper NE reassembly and can drive spontaneous NE rupture exposing nuclear DNA to the cytosol allowing unrestricted cGAS activation and STING-mediated apoptosis.

Interestingly, in humans, ANKLE2 is a target of Zika virus protein NS4A which antagonizes its activity ultimately leading to microcephaly^63^. Humans carrying homozygous or compound heterozygous mutations in either ZNF335 or ANKLE2 exhibit severe microcephaly like that characteristic of Zika patients^34, 64^. Recent studies have demonstrated a critical role for central nervous system immune cells in regulating neuronal stem cell maintenance and differentiation. Specifically, microglia play a key role in this process^65–67^. Under conditions which stimulate cGAS activity, microglia and other CNS immune cells preferentially undergo apoptosis^68^. Based on the mechanism revealed in this study it is possible that microcephaly resulting from Zika infection or loss of ZNF335 or ANKLE2 may be driven by cGAS/STING-dependent apoptosis of neuronal progenitors and/ or CNS immune cells. Should our mechanism extend to neuronal progenitors or CNS immune cells it may be possible to pharmaceutically prevent microcephaly in these specific instances by inhibition of the cGAS/STING pathway. However, further research will be required to determine the viability of such a therapeutic approach.

## Acknowledgements

This study was funded by the NIH (R01-GM059638 and P01-AI102853) to YZ and (P01-AI102853) to DW.

We thank M. Cook, N. Martin, B. Li and L. Martinek (Duke University Cancer Institute Flow Cytometry Core) for technical support and cell sorting. We thank the Duke Molecular Physiology Institute for preparation of scRNA-seq libraries. We thank M. Krangel, QJ Li, J. Racine, D. Serreze, and M. Hasham for critical reading and comments on the manuscript. We thank M. Ciofani and J. Park for providing cell lines and mice. We thank M. Parker and J. Wheaton of M. Ciofani’s lab for providing MSCV-Thy1.1 and MSCV-sgRNA expression vectors.

## Author Contributions

JJR, YZ and DW designed experiments and analyzed and interpreted data. JJR, QW, NM, DD, MJH, SW, SR and AVC performed experiments. JJR, YZ and DW wrote the manuscript with editing by the co-authors. JJR and YZ oversaw and supervised all aspects of the study.

## Methods

### Mice

B6.Cg-*Zfp335^tm1Caw^* (Zfp335^fl/fl^, Stock No. 022413) and B6J.129(B6N)-*Gt(ROSA)26Sor^tm1(CAG-cas9*,-EGFP)Fezh^/*J (R26^LSL-Cas9^, Stock No. 026175) mice were purchased from The Jackson Laboratory. C57BL/6J-Tg(Cd8a*-cre)B8Asin (E8III-cre) mice were generously provided by Jung-Hyun Park (NIH). B6.129S-Tcrd^tm1.1(cre/ERT2)Zhu^ (*Tcrd^CreERT2^*) have been maintained in our colony since original development. A modified Ai6 targeting vector to drive conditional overexpression of Bcl2 was generated by cloning in mouse *Bcl2* cDNA (Transomic Technologies) using FseI and SfiI restriction sites. R26^LSL-Bcl2^ mice were generated by the Duke University Transgenic Facility using G4 mouse embryonic stem cells. Animals were maintained under specific pathogen-free conditions at the Cancer Center Isolation Facility of Duke University Medical Center. All experimental procedures were approved by the Institutional Animal Care and Use Committee. All mice used in this study were 4-8 weeks old. For all experiments Cre-negative littermate controls were used unless otherwise stated.

### Antibodies

All antibodies used in this study were purchased commercially and have previously been validated. Anti-TCRγδ (GL3) was purchased from BD Biosciences. Anti-TCRγδ (GL3), rabbit anti-Lamin B (10H34L18), polyclonal rabbit anti-Banf1 (Cat. PA5-20329) and goat anti-rabbit IgG (H+L)-Alexa Fluor 647 were purchased from ThermoFisher Scientific. Anti-CD16/32 (2.4G2) was purchased from Tonbo Biosciences. Anti-CD90.1 (OX7), anti-CD90.2 (30-H12), anti-CD4 (RM4-5), anti-CD8 (53-6.7), anti-CD44 (IM7), anti-CD25 (PC61), anti-CD62L (MEL-14), anti-TCRβ (H57-597), anti-CD27 (LG.3A10), anti-Bcl2 (BCL/10C4), anti-CD24 (M1/69), anti-B220 (RA3-6B2), anti-CD11b (M1/70), anti-CD11c (N418), anti-CD19 (6D5), anti-Ly6G/Ly6C (RB6-8C5), anti-NK1.1 (PK136), anti-TER119 (TER-119), anti-CD117/c-kit (2B8), anti-Phosphoserine (M380B), mouse IgG1 isotype control (MG1-45), mouse IgG1 isotype control (MOPC-21) and Annexin V were purchased from Biolegend.

### Flow cytometry and cell sorting

Thymus or spleen tissues were harvested from 4-8 week old mice. Tissues were then dissociated in FACS Buffer (PBS supplemented with 2.5% FBS and 2mM EDTA) using a Dounce Homogenizer and filtered through 70µm nylon mesh (Genesee Scientific) to yield single-cell suspensions. For spleen samples, red blood cells were lysed using 1x RBC lysis buffer then resuspended in FACS buffer. 0.5-1x10^7^ cells were stained with fluorescently labelled antibodies for 30 minutes at 4°C then washed with excess FACS buffer. Prior to analysis propidium iodide (Sigma-Aldrich, Cat. P4170) or DAPI (Sigma-Aldrich, Cat. D9542) were added to a final concentration of 0.5µg/mL or 100ng/mL, respectively for live/ dead discrimination. Cells were analyzed on a Fortessa X20 (BD Biosciences) or FACSCantoII (BD Biosciences) cytometer. For isolation of thymocyte subsets or virally transduced cells, sorting was performed using a FACSDiva (BD Biosciences) or Astrios (Beckman-Coulter) cell sorter. For sorting of thymocyte subsets *ex vivo*, staining included a lineage dump stain consisting of B220, CD11b, CD11c, CD19, GR-1, NK1.1, TCRβ, TCRγδ and TER119 antibodies. All analyses were performed using FlowJo v10 software (TreeStar).

### Bulk RNA-seq

DP thymocytes (Lin^-^ CD4^+^ CD8^+^) were FACS sorted from total thymus of 7-week-old female Zfp335^fl/fl^ E8_III_-cre or Zfp335^+/+^ E8_III_-cre mice. Purified DP cells were lysed with Trizol and RNA isolated using the DirectZol Micro RNA prep kit (Zymo) according to manufacturer’s recommended protocol. gDNA was eliminated by on-column DNase digestion. Libraries were prepared using standard preparation protocols by BGI Genomics. 150bp paired-end sequencing was performed on the BGISEQ-500 sequencing platform.

Paired-end reads were mapped to the mouse mm10 reference genome using the HiSat2 software and count matrices generated using the featureCounts function of the Subreads software package. Differential expression analysis was performed using edgeR and DeSeq2 implemented through iDep.91 (http://bioinformatics.sdstate.edu/idep90/). Gene-Set Enrichment Analysis (GSEA) was utilized to identify enriched pathways based on differential expression analysis using pre-ranked gene lists.

### Cell Culture

OP9-DL1 cells, kindly provided by Maria Ciofani (Duke University) were cultured in MEMα (Gibco) supplemented with 10% FBS (Atlanta Biologicals) and 1x penicillin/ streptomycin (Gibco). HEK293T cells were cultured in DMEM supplemented with 10% FBS, 1x penicillin/ streptomycin, 1x non-essential amino acids and 1x GlutaMAX. For OP9-DL1 culture of thymocytes, cultures were additionally supplemented with 5ng/mL recombinant mouse IL-7 (Biolegend). Scid.adh.2c2 cells were cultured in IMDM supplemented with 10% FBS (Hyclone), 1x penicillin/ streptomycin, 1x NEAA, 1x sodium pyruvate, 1x GlutaMAX, and 55µM β-mercaptoethanol. In some OP9-DL1 cultures 5µg/mL RU.521 (Invivogen), 0.5µg/mL H-151 (Cayman Chemicals) or 25µg/mL Cridanimod (Cayman Chemicals) were added. All cultures were maintained at 37⁰C with 5% CO_2_.

### DN thymocyte enrichment

Total thymocytes were harvested from 4–8-week-old mice. Tissues were dissociated and strained through 30µm nylon mesh (Genesee Scientific). For purification of DN3/4 thymocytes cells were stained with biotinylated antibodies against B220, CD3, CD4, CD8, CD11b, CD11c, CD19, CD44, c-Kit, GR-1, IgM, NK1.1, TCRβ, and TCRγδ. For enrichment of total DN cells CD44 and c-Kit antibodies were excluded. Following antibody staining, cells were incubated with 50µL or 100µL of streptavidin magnetic particles (Spherotech, cat. SVM-40-100) / 10^7^ cells at 2 x 10^7^ cells/mL in FACS buffer for total DN enrichment or DN3/4 purification, respectively. Particle-bound cells were separated three times on a magnetic rack.

### Retrovirus packaging and transduction

Retrovirus were generated by transfecting HEK293T cells with 1µg/mL each of MSCV transfer and pCL-Eco vectors using Lipofectamine 2000 (Invitrogen) or JetOptimus (Genesee Scientific) according to manufacturer’s recommended protocols. Media was changed 24 hours post-transfection and viral supernatants harvested 24 hours later. DN3/4-enriched thymocytes were transduced with fresh viral supernatant via spinfection for 2 hours at 2300 rpm at 30°C with 6.7µg/mL polybrene (Millipore). Following spinfection cells were transferred to culture on OP9-DL1 stromal cells for overnight culture. 18-24 hours post-infection virally transduced (DsRed+ or Thy1.1+) DN3 (CD25+) or DN4 (CD25-) were isolated by FACS sorting for an additional 3-5 days of culture in the OP9-DL1 culture system. For dual-targeting CRISPR experiments, equal volumes of sgRNA-Thy1.1 and -DsRed viral supernatants were mixed for transduction.

### scRNA-seq library preparation

For single cell RNA-sequencing, DN4 thymocytes (Live Lin^-^ CD4^-^ CD8^-^ CD25^-^ CD44^-^) were sorted from one male and one female mouse pooled for each genotype using an Astrios Sorter. Sorted cells were encapsulated into droplets and libraries were prepared using a Chromium Single Cell 3’ Kit using the v3.1 chemistry. 7,000 cells per genotype were targeted. scRNA-seq libraries were pooled and sequenced on a NovaSeq S Prime Flow Cell yielding an average depth of 71,584 or 67,816 reads per cells for Zfp335cKO or WT samples, respectively.

### scRNA-seq analysis

scRNA-seq data were processed using the Cell Ranger pipeline (10x Genomics). FASTQ files were generated from raw base call logs (bcl2fastq, v2.20), aligned to the mouse mm10 (release 93) reference genome (cellranger, v3.1.0; STAR v2.5.3a) to generate raw gene count matrices.

All downstream analyses were performed using the R software package Seurat (v4.0.0). Data was filtered to exclude cells with < 1,000 genes detected or < 1,000 UMIs. Doublets were excluded by filtering cells with > 60,000 UMIs. Low-quality cells were further filtered by removal of cells with > 7.5% mitochondrial gene expression. Gene expression matrices were then merged, data normalized, scaled and cell cycle scored using standard methods with Seurat. Dropouts were imputed using the R package ALRA. Cell cycle phase was regressed, and principal component analysis (PCA) was performed on the 6,000 most variable genes. 35 principal components were selected for downstream analysis based on JackStraw analysis. Dimensionality reduction was performed by Uniform Manifold Approximation and Projection (UMAP) and clustering defined using a resolution of 0.5. Gene expression was visualized by VlnPlot, DotPlot and FeaturePlot functions in Seurat. Gene signature scores were calculated using SingleCellSignatureExplorer and previously described methods ^69^. Differential expression analysis was performed using the FindMarkers function in Seurat with Wilcoxon Rank Sum Test.

### Cloning cDNA overexpression vectors

Bcl2 overexpression vector was generated by cloning Bcl2 cDNA (Transomic Technologies, Cat. TCM1304) into the pMSCV-loxp-dsRed-loxP-eGFP-puro-WPRE vector (Addgene #32702) using the EcoRI and NsiI restriction sites. Ankle2 cDNA (Transomic Technologies, Cat. TCM1004) was cloned into the MSCV-IRES-Thy1.1 vector using NEBuilder Hifi Assembly (New England Biolabs). All vectors were propagated in Stbl3 cells (ThermoFisher Scientific).

Generation of *Scid.adh.2c2-dCas9^10x-GCN4^* CRISPRi cells dCas9^10x-GCN4^ (pHRdSV40-dCas9-10xGCN4_v4-P2A-BFP, Addgene #60904) was lentivirally transduced into Scid.adh.2c2 cells, following which BFP+ cells were isolated by flow cytometry. Single cells were then cloned into 96 well plates and screened for knockdown efficiency using CD25 gRNA retroviral vectors. Clones exhibiting more than 90% CD25 downmodulation were expanded for use in our studies.

### Generation of gRNA retroviral vectors

All gRNAs were designed using the CRISPick ^70^ gRNA design tool. All gRNAs were cloned into expression vectors by annealing followed by ligation into a BbsI cleavage site. The basic gRNA expression vector used was the MSCV-mU6-sgRNA-hPGK-Thy1.1 (kindly provided by Maria Ciofani). Knock-out gRNAs were first cloned into this Thy1.1 backbone. To generate DsRed expressing vectors for dual targeting, Thy1.1 was removed by digestion with BamHI and EcoRI and replaced with DsRed Express II by NEBuilder Hifi Assembly. The CRISRPi retroviral vector was generated by first cloning the pSV40-scFv-GCN4-sfGFP-VP64-GB1-NLS (Addgene #60904) fusion construct into the MSCV-mU6-sgRNA-hPGK backbone followed by replacement of VP64 with KRAB using NEBuilder.

### qPCR analysis of gene expression

Following viral transduction, Scid.adh.2c2.dCas9^10x-GCN4^ cells were assessed for transduction efficiency by flow cytometry. For samples exceeding 90% GFP+ 10^6^ cells were lysed in Trizol and RNA isolated using the Direct-Zol MicroPrep kit. 500ng of RNA was reverse transcribed using SuperScript III Reverse Transcriptase (Invitrogen) with random hexamers according to the manufacturer’s recommended protocol. 5ng of cDNA per 25µL reaction was then used for gene expression analysis with PowerTrack Sybr Green Master Mix (Applied Biosciences) according to the manufacturer’s recommended protocol using fast cycling conditions with an Eppendorf MasterCycler qPCR machine. Relative expression was determined using the ddCt method with Gapdh being used for normalization.

### Proximity Ligation Assay for Baf phosphorylation

Proximity ligation assays were performed using the Duolink^®^ flowPLA Detection Kit – Orange (Millipore Sigma, Cat. DUO94003) according to manufacturers recommended protocol with minor changes. Briefly, total thymocytes were prepared as described in flow cytometry and cell sorting methods section. 10^7^ thymocytes were stained with surface antibodies to distinguish all major thymocyte subsets. Next, cells were fixed with 4% paraformaldehyde for 10 minutes, washed and permeabilized for 30 minutes at room temperature. After permeabilization, cells were blocked for 1 hour at 37⁰C with 300µl of Duolink^®^ blocking solution then stained overnight at 4⁰C with purified mouse anti-Phosphoserine (Biolegend) or purified mouse IgG1 isotype control (clone MG1-45, Biolegend) and purified rabbit anti-Banf1 (ThermoFisher Scientific) or rabbit anti-Lamin B1 (ThermoFisher Scientific) diluted in Duolink^®^ antibody diluent. After each step cells were washed twice with 1mL or Duolink^®^ In Situ wash buffer. Next, cells were incubated for 1 hour at 37⁰C with Duolink^®^ In Situ PLA^®^ Probe Anti-Rabbit PLUS (Sigma Millipore, Cat. DUO92002) and Duolink^®^ In Situ PLA^®^ Probe Anti-Mouse MINUS (Sigma Millipore, Cat. DUO92004) diluted in Duolink^®^ antibody diluent. Additional controls in which individual probes were omitted were also prepared. Following probe incubation, cells were washed then incubated for 30 minutes at 37⁰C with 1x Duolink^®^ ligation reaction mixture, washed again and incubated for 90 minutes with 1x Duolink^®^ amplification reaction mixture. Following amplification, cells were incubated with 1x Duolink^®^ Detection Solution – Orange for 15 minutes at 37⁰C. Cells were finally washed, resuspended in PBS and assayed using a FortessaX20 cytometer (BD Biosciences).

### Determination of nuclear envelope structure

5x10^4^ Hela cells per well were reverse transfected with 15pmol siRNA using Lipofectamine RNAiMax (ThermoFisher Scientific) in an 8 well chamber slide according to recommended protocols. ANKLE2 and universal non-targeting control siRNAs were purchased from IDT (Design ID: hs.Ri.ANKLE2.13). BANF-targeting siRNAs were purchased from ThermoFisher Scientific (IDs: s16807, s16808, 26065). 48 hours post-transfection cells were fixed with 4% paraformaldehyde for 10 minutes at room temperature and permeabilized with permeabilization buffer for 1h at RT temperature. Primary antibody Lamin B (Invitrogen, Cat. 702972) were added for overnight incubation at 4C and washed with 1X PBS for three time. After that, secondary antibody Alexa Fluor 647-conjugated goat anti-rabbit antibody (Invitrogen, Cat. A32733) were added for 12h at 4C in the dark. After washing with 1X PBS for three times, slides were mounted with DAPI-containing mounting media (VECTORLAB, Cat. H-1200). Images were collected using Zeiss 780 upright confocal.

To analyze nuclear structure DAPI channel images were converted to binary with ImageJ. Following binarization, the Watershed function was used to separate touching cells. Circularity was then determined with a minimum threshold of 500 px^2^.

### *In vivo* H-151 treatment of mice

Mice were administered 750 pmol (210µg) of H-151 (Cayman Chemicals) or vehicle via intraperitoneal injection daily for 7 days beginning at 7 weeks of age. The vehicle for injections was sterile PBS + 10% Tween-80 (VWR).

### Statistical analysis

Statistical tests were performed using GraphPad v9.0.0 (Prism). For graphs with multiple comparisons being made, two-way ANOVA was performed with post-hoc Sidak’s test or Tukey’s test for multiple comparisons. For comparisons of cell numbers, data was log transformed prior to statistical tests. For all Two-way ANOVA tests normality tests were performed to ensure normalcy assumptions were met. For graphs of single comparisons, a two-tailed Mann-Whitney test was used. All significant p-values are shown in each graph. No statistical methods were used to predetermine sample size.

### Data and code availability

Data generated in this study can be accessed upon publication through NCBI Gene Expression Omnibus (https://www.ncbi.nlm.nih.gov/geo/) under accession GSE189244.

## Supplementary Information

**Supplementary Figure 1.**
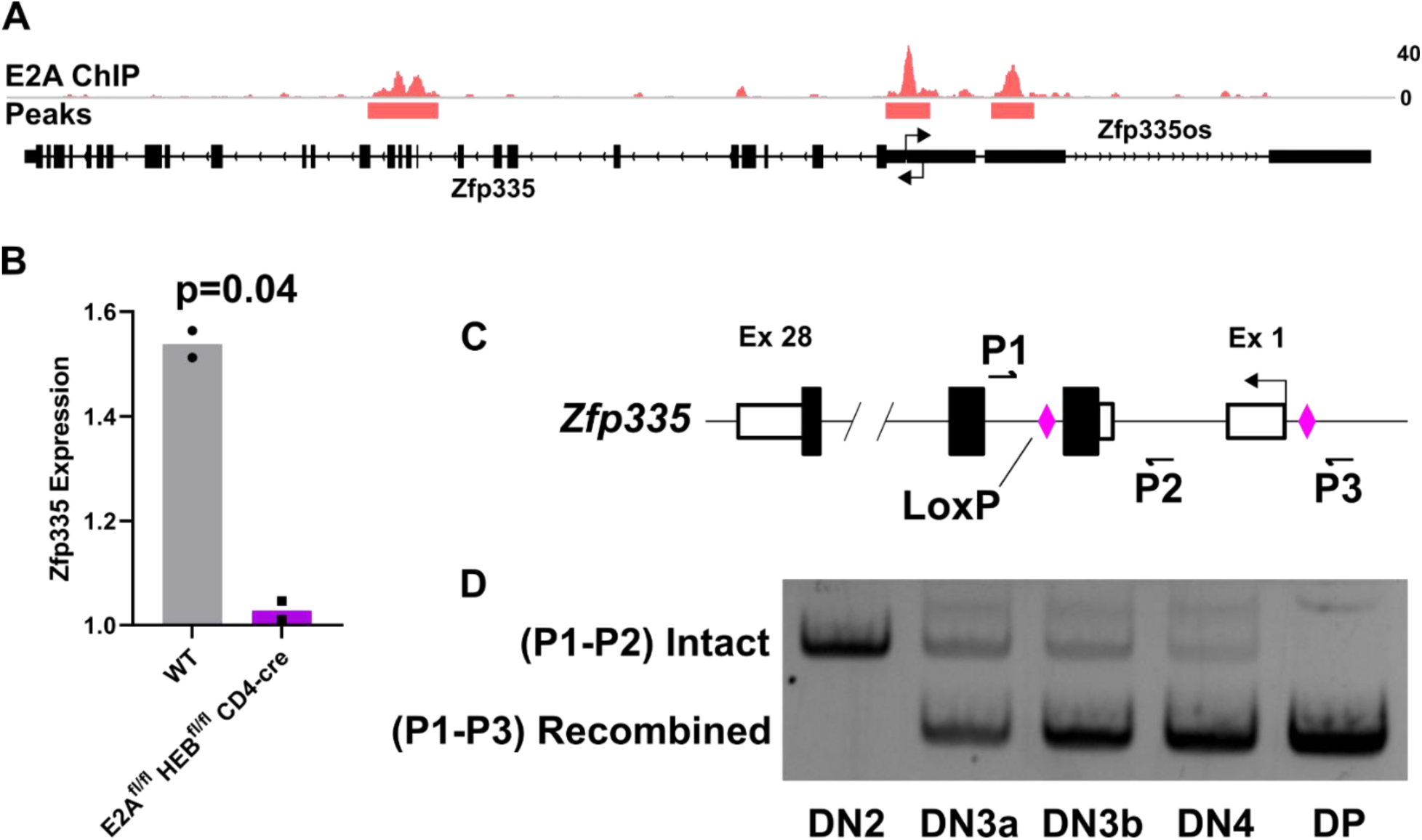
Zfp335 is a target of E proteins in developing T cells. (A) E2A ChIP-seq track for *Zfp335* locus in *Id2^fl/fl^ Id3^fl/fl^ Lck-cre* DP thymocytes (GSE89849). (B) Zfp335 transcript abundance in WT vs. E2A/HEB double knock-out DP thymocytes determine by microarray (GSE9749). (C) Schematic diagram for PCR-based determination of Zfp335 recombination kinetics. Small arrows indicate approximate positions for primers (P1-3) used for assay. (D) Representative assessment of Zfp335 recombination in sort purified *Zfp335^fl/fl^ E8III-cre* DN2, DN3a, DN3b, DN4 or DP thymocytes. Data are representative of four individual experiments.

**Supplementary Figure 2.**
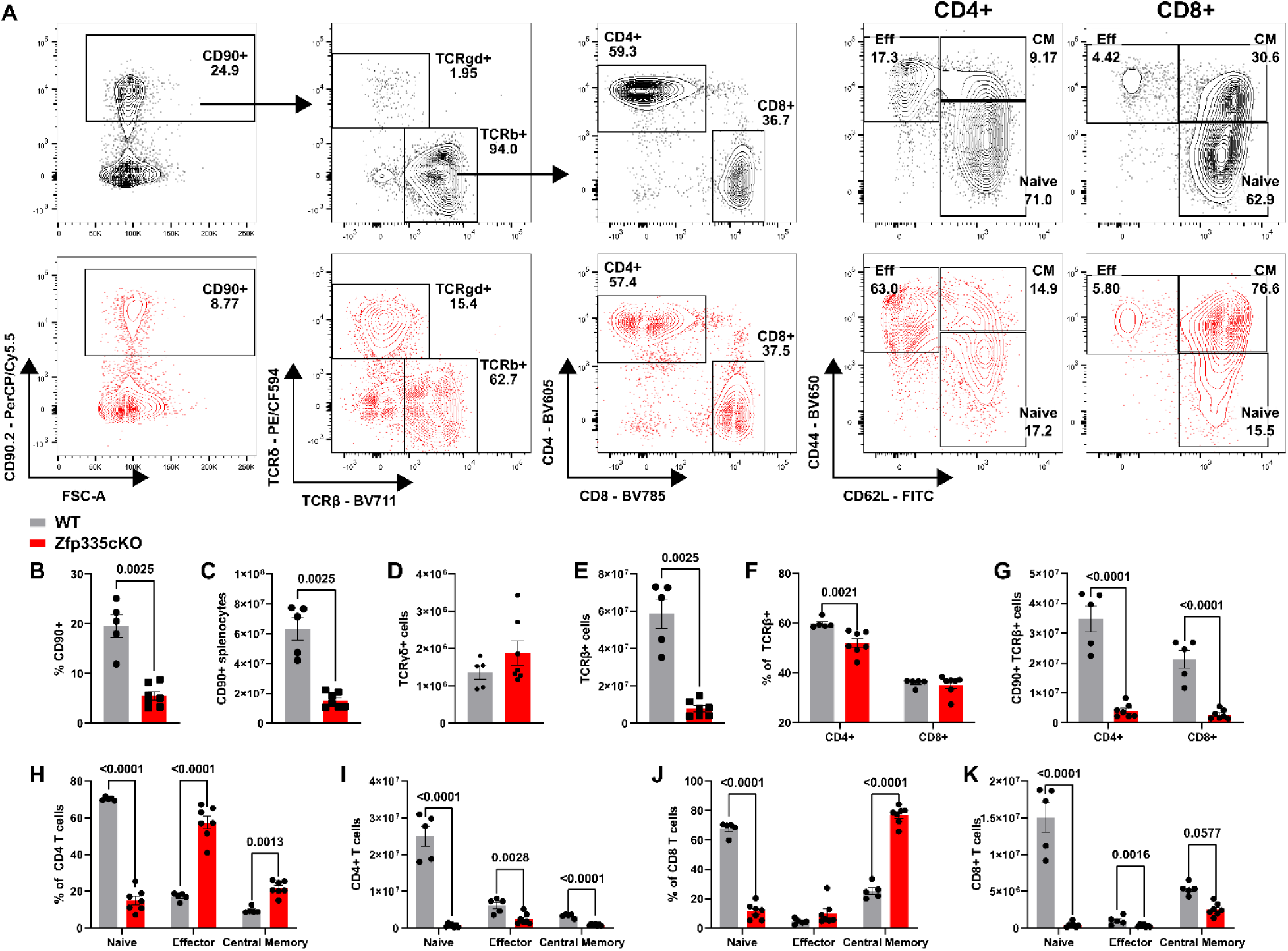
Zfp335cKO mice exhibit T lymphopenia and reduced peripheral naïve T cells. (A) Gating schema for identification of WT (black) or Zfp335cKO (red) splenic T cell populations beginning with live (DAPI^-^) splenocytes. Proportion (B) or total numbers (C) of splenic CD90+ cells. Total numbers of TCRγδ+ (D) or TCRαβ+ (E). Proportions (F) and total numbers of CD4+ or CD8+ TCRαβ cells. Proportions (H, J) and numbers (I, K) of naïve, effector or central memory T cells within the CD4+ or CD8+ compartment. WT (n=5) or Zfp335cKO (n=7) from two separate experiments. *P*-values determined by Mann-Whitney U-test (B-E) or Two-Way ANOVA with *post hoc* Sidak test (F-K). Plots show mean ± sem.

**Supplementary Figure 3.**
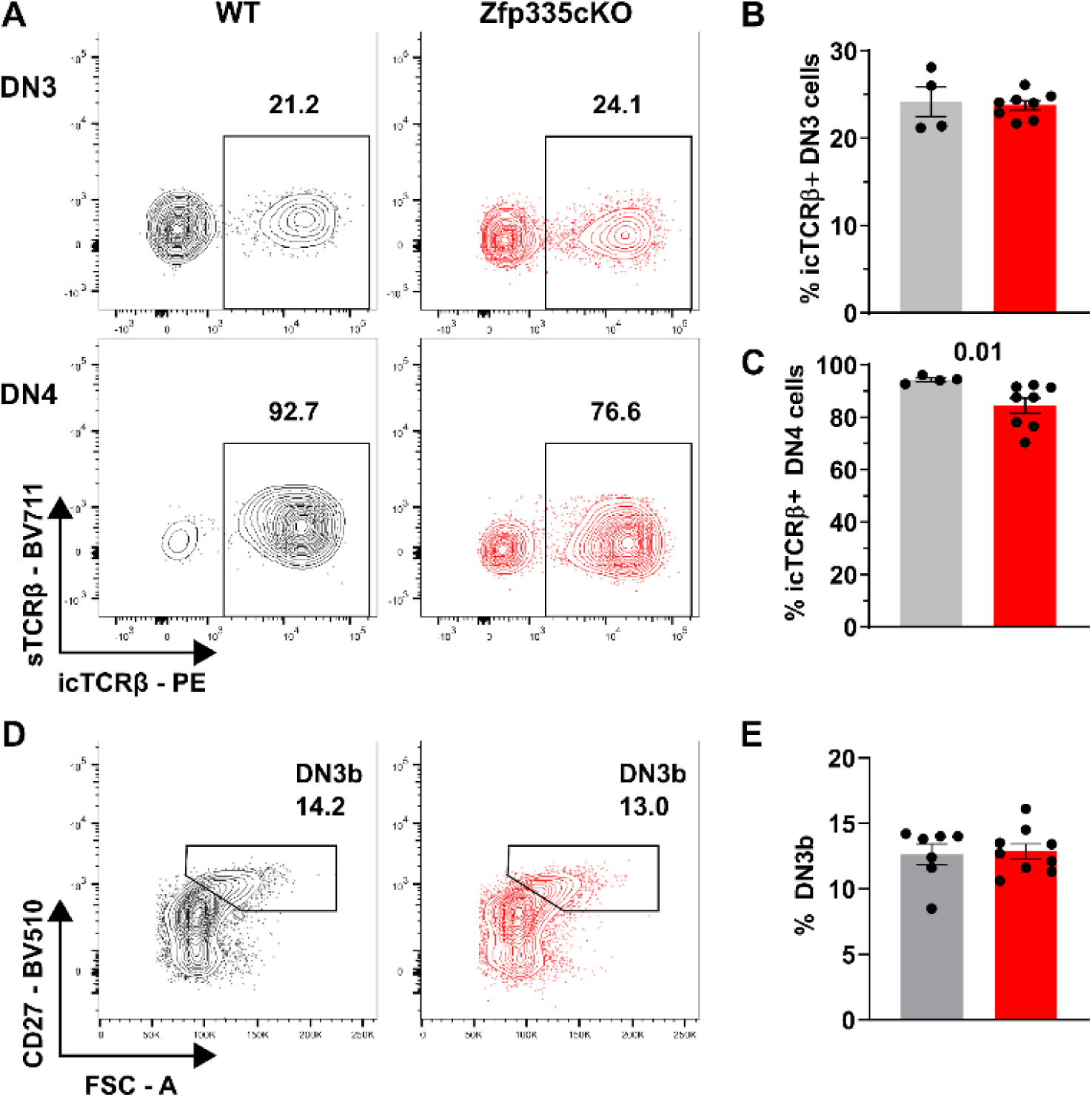
Loss of Zfp335 during DN3 does not impair β-selection. (A) Gating for icTCRβ expression among DN3 (CD90^+^ TCRδ^-^ CD4^-^ CD8^-^ sTCRβ^-^ CD44^-^ CD25^+^) or DN4 (CD90^+^ TCRδ^-^ CD4^-^ CD8^-^ sTCRβ^-^ CD44^-^ CD25^-^) thymocytes. Frequency of icTCRβ DN3 (B) or DN4 (C) cells among WT (n=4) or Zfp335cKO (n=8) mice. (D) Flow cytometric gating for identification of WT or Zfp335cKO DN3b thymocytes pre-gated on total DN3 cells. (E) Quantification of DN3b frequency among WT or Zfp335cKO DN3 thymocytes. *P-*values determined by Two-way ANOVA with *post hoc* Sidak test (B,C) or Mann-Whitney U-Test (E). Plots show mean ± sem.

**Supplementary Figure 4.**
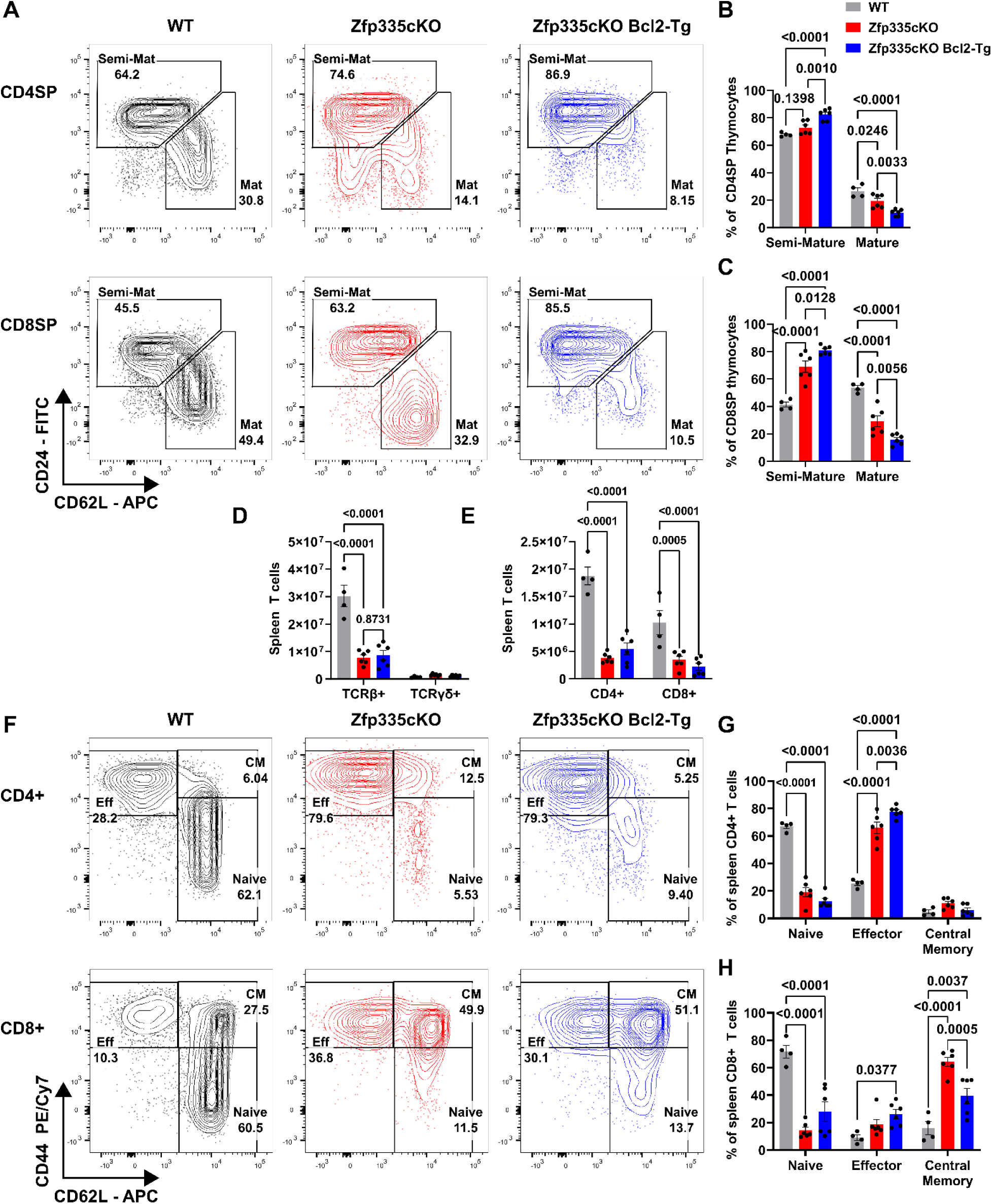
Bcl2 overexpression fails to rescue thymic differentiation defect and peripheral T lymphopenia in Zfp335-deficient mice. Representative gating (A) and quantification of CD4SP (B) or CD8SP (C) thymic maturation. Total splenic TCRβ and TCRγδ (D) T cells. Quantification of total splenic CD4+ or CD8+ TCRβ+ T cells. Representative gating (F) and quantification of splenic CD4+ (G) or CD8+ T cell effector status. n=4 WT, n=6 Zfp335cKO, n=6 Zfp335cKO Bcl2-Tg. Data are compiled from three independent experiments. *P-*values determined by Two-Way ANOVA with *post hoc* Sidak Test. Plots show mean ± sem.

**Supplementary Figure 5.**
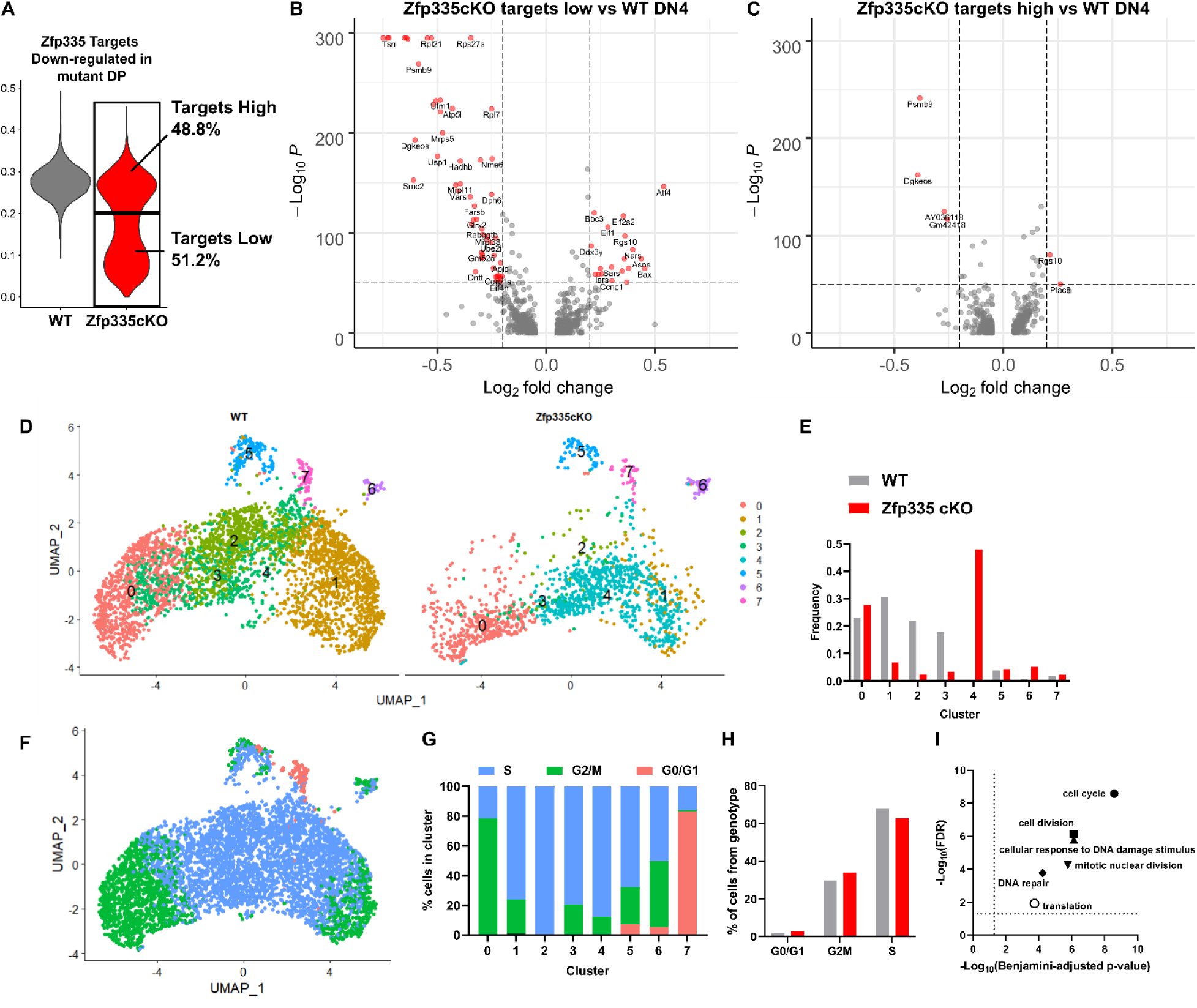
scRNA-seq identifies ‘true’ Zfp335 mutant DN4 cells. (A) Violin plot of gene set score for Zfp335 target genes down-regulated in mutant DP thymocytes (Fig 1L-M) and cutoff value used to identify ‘true’ Zfp335 mutant cells with low target score (lower box) and non-mutant cells (upper box). Volcano plots of differentially expressed genes between Zfp335cKO targets low (B) or Zfp335cKO targets high (C) cells compared with WT control. (D) UMAP projections colored by cluster and separated by genotype for WT and true Zfp335 mutant DN4 cells. (E) Frequency of cells found within each cluster. UMAP projection (F) and quantification of cell cycle phase for each cluster (G). (H) Quantification of distribution of cell cycle phase by genotype. (I) GO analysis of top 25 cluster defining genes for each cluster (dashed lines indicate significance cutoff of p<0.05 and FDR<0.05. *P*-values determined by Wilcoxon Rank Sum test (B-C) or Fischer’s Exact test with Benjamini-Hochenberg correction (I).

**Supplemental Figure 6.**
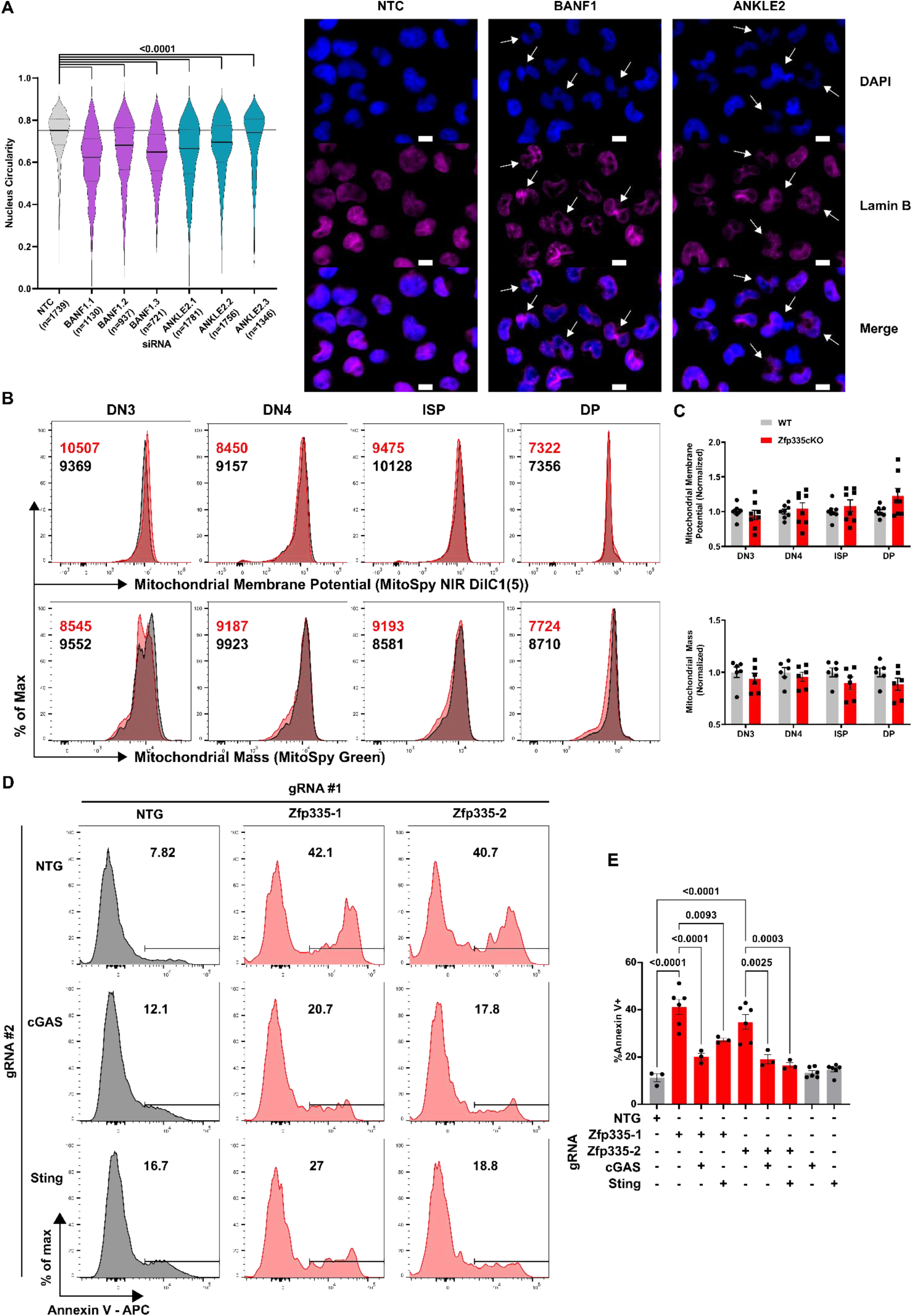
Ankle2/BANF1 control nuclear envelope architecture. Quantification of nuclear circularity (A) in Hela cells transfected with non-targeting control (NTC), BANF1-, or ANKLE2-targeting siRNAs 48 hours post-transfection (left). Representative images DAPI or Lamin B staining of siRNA transfected Hela cells (right). Scale bars are 10µm. Arrows indicate cells with severely disrupted nuclear envelope architecture. Representative histograms (B) and compiled data (C) for mitochondrial membrane potential (top) or total mitochondrial mass (bottom) in WT or Zfp335cKO thymocyte populations *ex vivo*. Representative gating (D) and quantification (E) of Annexin V binding among DN4 cells from *R26^LSL-Cas9^ E8III-cre* DN3/4 thymocytes transduced with indicated gRNA-expressing retroviruses and cultured for three days on OP9-DL1 cells. P-values calculated using One-Way ANOVA with Dunnett’s post hoc test. Plots show mean ± sem (G) or median (solid line) and interquartile range (dotted lines) (E). Data are compiled from two (B-E) or three (F-G) independent experiments.

**Table S1.**
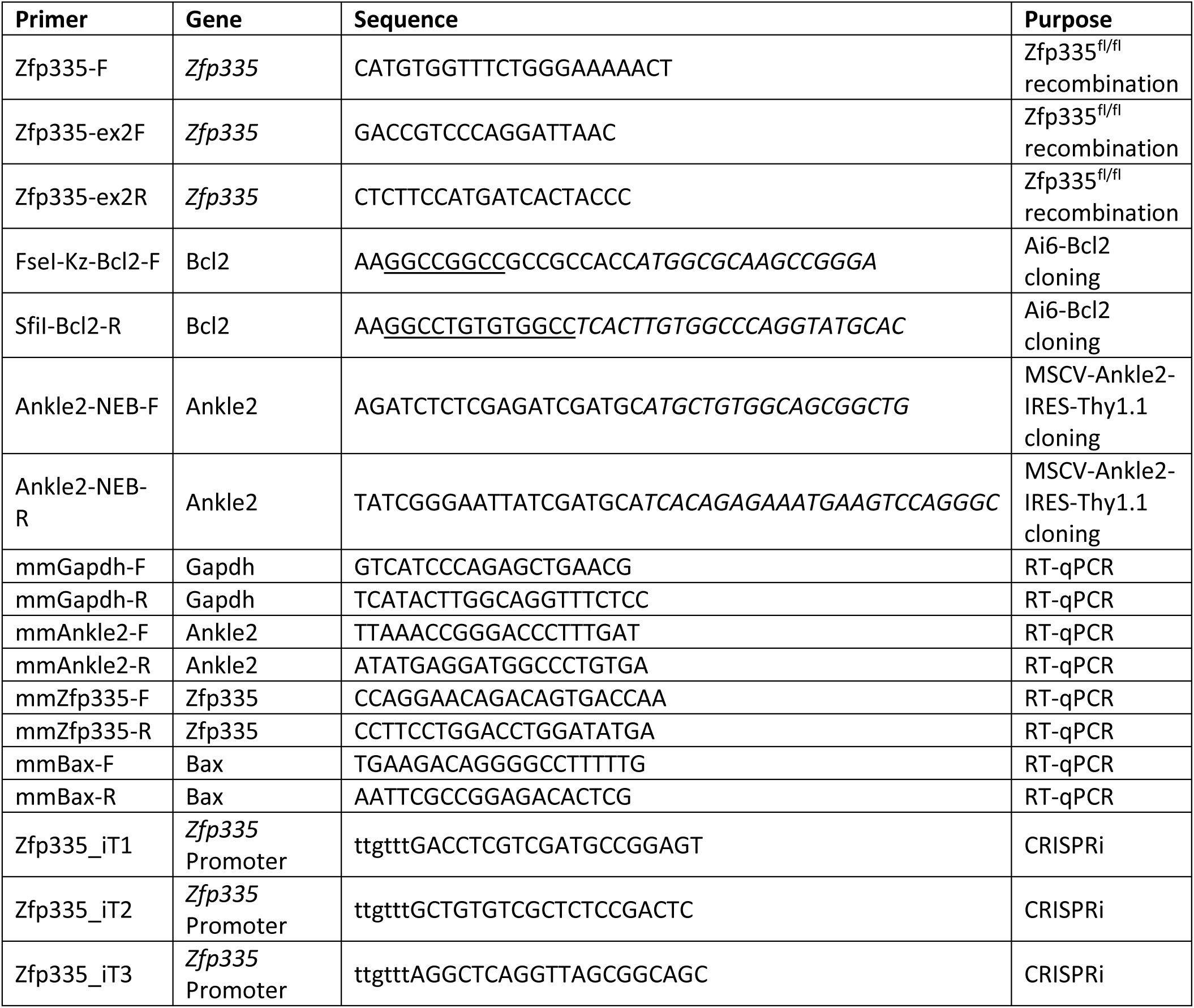
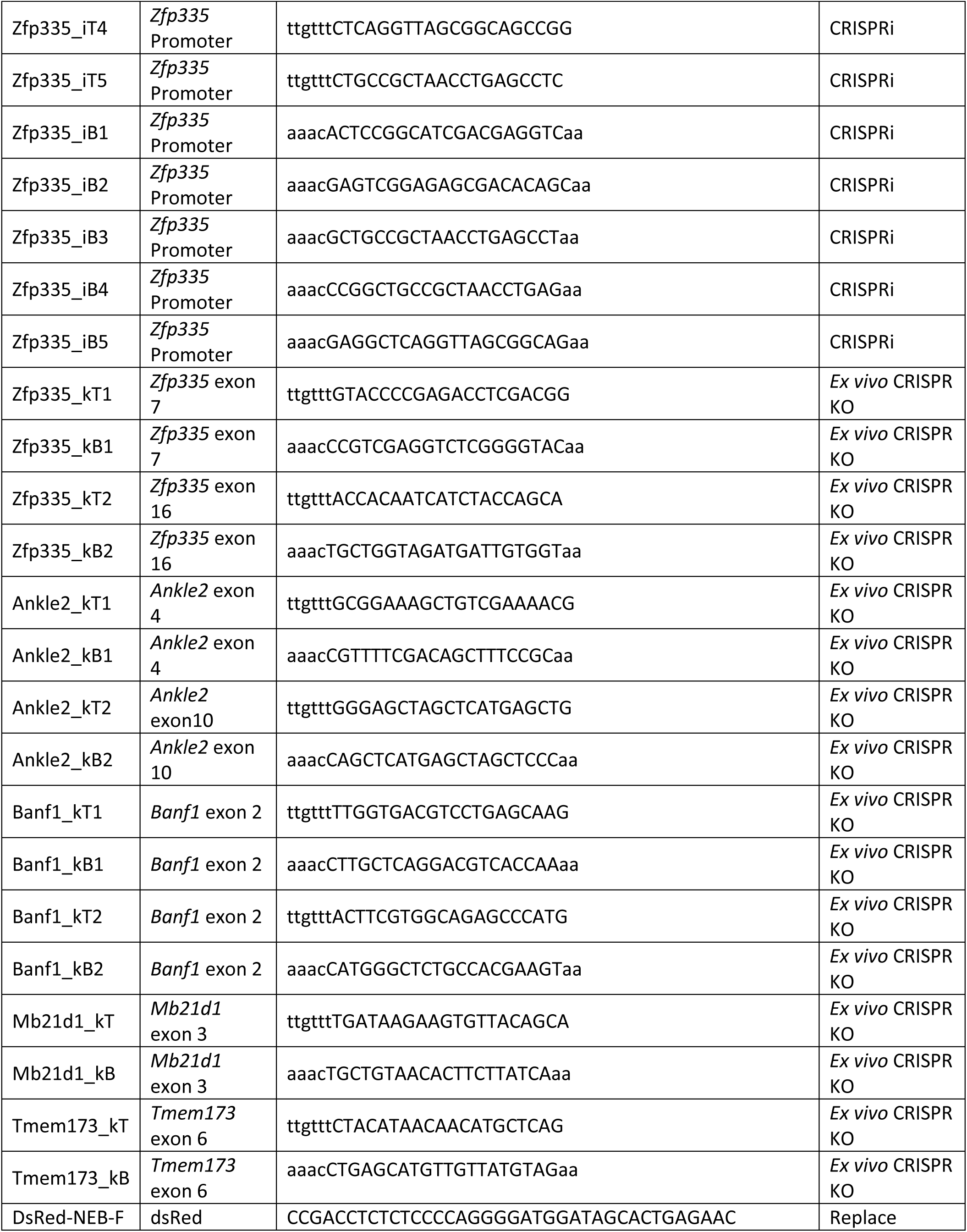
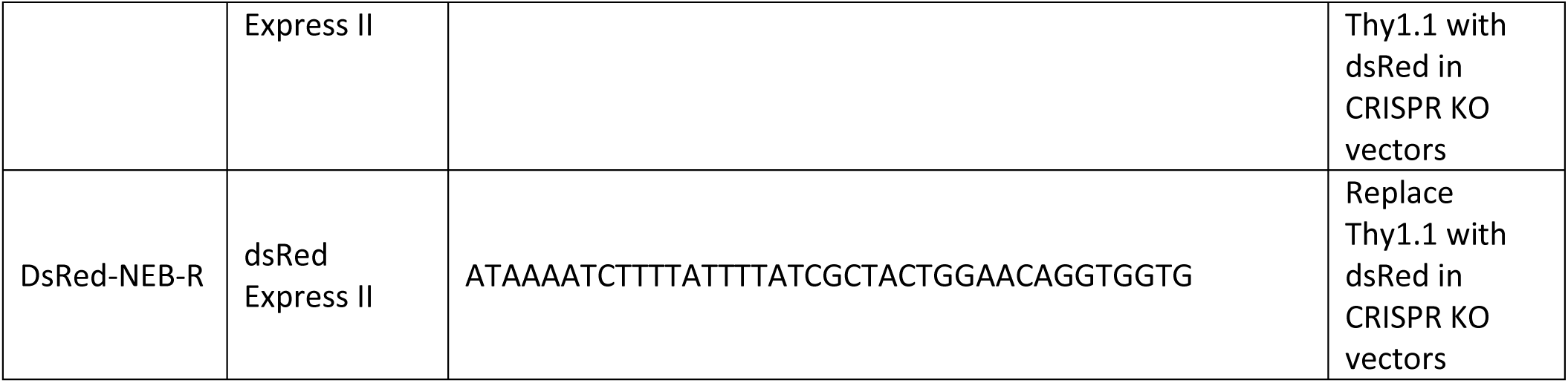
Primer sequences (Related to Figures S1, 3, 5, 6 and S6)

